# Metabolic immunity to infection is driven by mitochondrial one-carbon metabolism

**DOI:** 10.1101/2025.05.07.652698

**Authors:** Tânia Catarina Medeiros, Silvia Reato, Xianhe Li, Bruna Martins Garcia, Irina Rais, Kira Allmeroth, Matías D. Hartman, Martin S. Denzel, Martin Purrio, Andrea Mesaros, Kit-Yi Leung, Nicholas D.E. Greene, Patrick Giavalisco, Lena Pernas

**Affiliations:** Metabolism of Infection Group, Max Planck Institute for Biology of Ageing, Cologne, Germany; Dept. Microbiology, Immunology & Molecular Genetics, University of California Los Angeles, USA; Howard Hughes Medical Institute, Chevy Chase, MD USA; Metabolic and Genetic Regulation of Ageing, Max Planck Institute for Biology of Ageing, Cologne, Germany; Phenotyping Core Facility, Max Planck Institute for Biology of Ageing, Cologne, Germany; Developmental Biology & Cancer Department, UCL, Great Ormond Street Institute of Child Health, University College London, UK; Metabolomics Core Facility, Max Planck Institute for Biology of Ageing, Cologne, Germany

## Abstract

As large consumers of cellular metabolites, mitochondria are positioned to compete with invading microbes for the nutrients they require to grow. Yet, little is known of whether cells weaponize mitochondrial metabolism during infection. We found that the transcription factor ATF4 activated a mitochondrial metabolic defense based on the essential B vitamin folate. During infection with the human parasite *Toxoplasma gondii*, ATF4 increased mitochondrial DNA (mtDNA) levels by driving the one-carbon (1C) metabolism processes that occur in mitochondria and use folate. The activation of ATF4 depended on host detection of parasite effector proteins, and restricted parasite growth by limiting its access to folate(s) required for dTMP synthesis. Impairing host mitochondrial 1C metabolism downstream of ATF4 promoted parasite growth, while forcing host dependence on mitochondrial 1C metabolism had the opposite effect. ATF4 activation also promoted a host-protective response in a mouse model of *Toxoplasma* infection. Thus, ATF4 rewires mitochondrial metabolism to activate a folate-based metabolic immunity against *Toxoplasma*. Our work paves the way for future studies exploring noncanonical defense strategies mediated by mitochondria and the role of folate metabolism during infectious disease.

**One-Sentence Summary:** ATF4 rewires mitochondrial metabolism during infection to drive a host-protective response based on folate competition.

## Main Text

A mammalian cell can contain hundreds to up to thousands of mitochondria (*1*). In a healthy cell, mitochondria are viewed as powerhouses that produce ATP and anabolic precursors, and signaling hubs that regulate diverse cellular programs (*2*). To carry out these functions, these organelles consume vast amounts of cellular nutrients and are equipped with a high density of porins and transporters that enable the uptake and release of metabolites (*3*). In an infected cell however, mitochondria are perceived as targets for intracellular microbes. This perception gives little importance to the fact that mitochondria can function as nutrient competitors during infection, thereby depriving intracellular pathogens access to key nutrients required for growth (*4, 5*). Yet, we do not know the extent to which host cells weaponize mitochondria in defense against pathogens, nor the mechanisms that cells use to activate mitochondrial metabolism for defense programs.

To investigate the cellular programs that rewire mitochondrial metabolism during infection, we focused on mitochondrial DNA (mtDNA). Unlike other organelles, mitochondria have their own genome, a vestige of their bacterial origin (*6*). To replicate their genome, mitochondria depend on cytosolic resources such as those sustained by the B vitamin folate (*7*). This essential cofactor is required for a broad set of reactions known as one-carbon (1C) metabolism that occur in the cytosol and mitochondria, and are crucial for many biochemical processes including the synthesis of purines and thymidine monophosphate (dTMP), both of which are used to generate deoxynucleotide triphosphates (dNTPs) for DNA replication (*7*). Thus, mtDNA levels reflect mitochondrial use of various cellular resources. It therefore comes as no surprise that dysregulated 1C metabolism and aberrant dNTP pools are linked to mtDNA replication defects associated with aging and neurodegenerative disease (*8, 9*).

As is the case for mitochondria, several intracellular pathogens require host products downstream of 1C metabolism for their growth. Host cells have therefore evolved defense mechanisms to limit their availability, such as through the degradation of dNTPs that occurs during viral infection (*5, 10*). However, whether cells deploy mitochondria to restrict pathogen access to 1C metabolism-linked nutrients is unknown. We used the protozoan pathogen *Toxoplasma gondii,* which infects up to one-third of the human population and has an unparalleled host range, to investigate the dynamics of mitochondrial metabolism during infection. *Toxoplasma* is of particular interest because like mitochondria, it salvages host resources that depend on 1C metabolism (*11, 12*).

### ATF4 regulates mtDNA levels during infection and infection-independent stress

To determine whether infection leads to changes in host mtDNA copy number, we measured mtDNA levels in human ES-2 ovarian cancer cells infected with *Toxoplasma.* We examined levels of *DLOOP*, the regulatory region of mtDNA, and 4 genes at different locations on the mitochondrial genome: *mt-ND1*, *mt-ND6*, *mt-CYTB*, and *mt-CO1* (Fig. 1A). At 24 hours post infection (hpi), we found that the copy number of each gene was significantly increased by ∼30% (Fig. 1B). Similar results were obtained in HeLa cells and human fibroblasts (HFF) (Supp. Fig. 1). *Toxoplasma* infection did not however lead to the release of mtDNA into the cytosol, as occurs during viral infections (Supp. Fig. 2) (*13*). Thus, *Toxoplasma* infection drives an increase in intramitochondrial DNA levels.

**Fig. 1.**
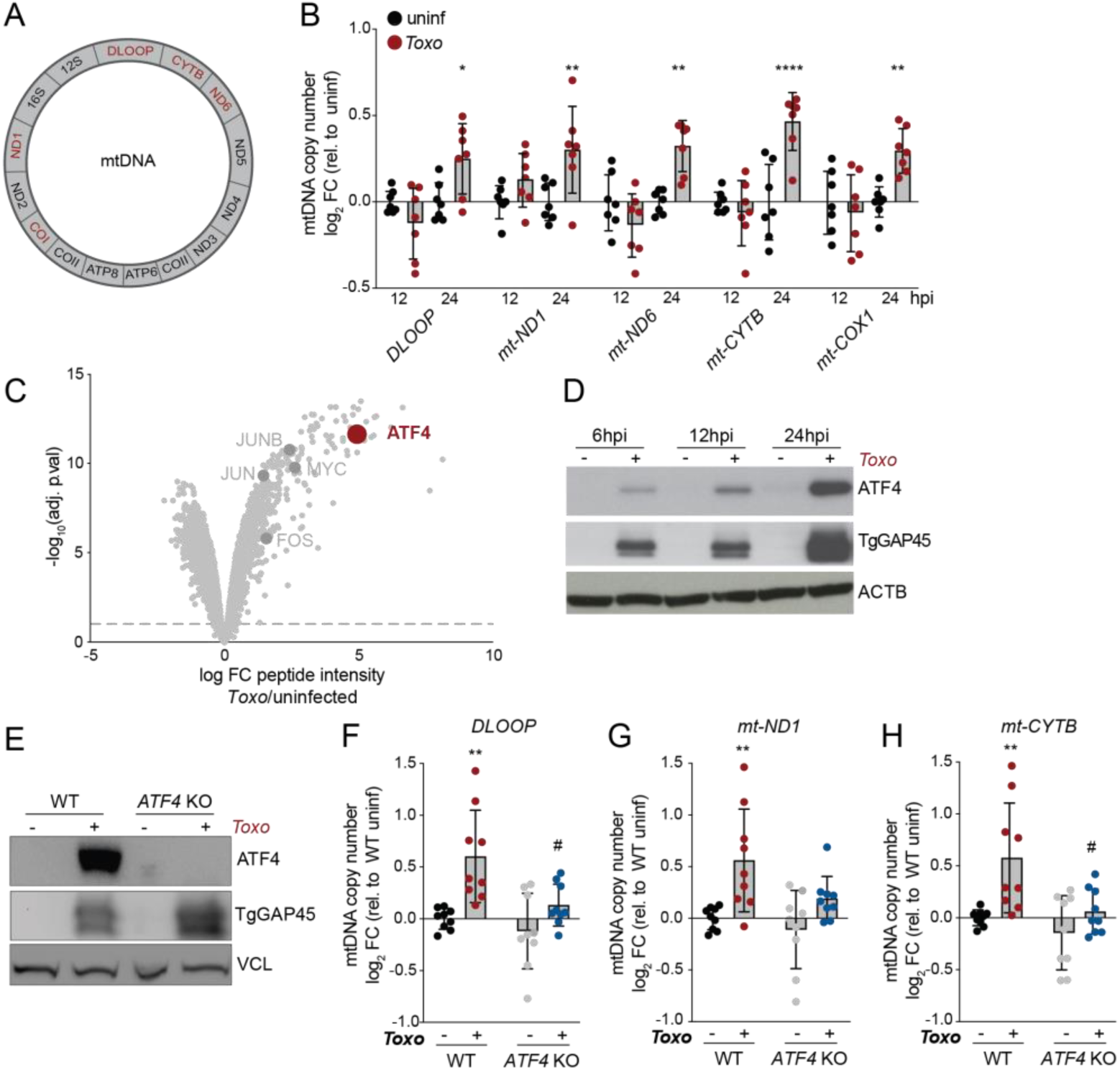
ATF4 regulates mtDNA levels during *Toxoplasma* infection. (**A**) Schematic of human mitochondrial genome, genes analyzed by qPCR in red. (**B**) mtDNA levels monitored by qPCR (normalized to *RUNX2*) in ES-2s cells that were uninfected or infected with *Toxoplasma* at a multiplicity of infection (MOI) of 4 at 12 and 24 hours post infection (hpi). Data are mean ± SD of n=7 independent cultures; *p<0.05; **p<0.01; ****p<0.0001 for uninfected versus infected by means of two-way ANOVA analysis. (**C**) Volcano plot of whole cell lysates from uninfected and infected HeLa cells at 24 hpi and analyzed by means of mass spectrometry with highlighted transcription factors. (**D**) ES-2 cells were uninfected or *Toxo-*infected (MOI: 4) for 6, 12, and 24 hours and analyzed by means of immunoblotting for ATF4, ∼50 kDa; β-Actin (ACTB), ∼45 kDa; and *Toxoplasma* GAP45 (TgGAP45), ∼45 kDa. (**E**) Immunoblot analyses of lysates from uninfected and *Toxoplasma-*infected (MOI: 4) WT and *ATF4* KO ES-2 cells at 24hpi: ATF4, ∼50 kDa; Vinculin (VCL), ∼124 kDa; and *Toxoplasma* GAP45 (TgGap45), ∼45 kDa. (**F, G** and **H**) mtDNA levels monitored by qPCR of *DLOOP*, mt-*ND1* and mt-*CYTB* (normalized to *RUNX2*) in uninfected and *Toxoplasma-*infected (MOI: 4) ES-2 cells at 12 at 24 hpi. Data are mean ± SD of n=9 independent cultures; **p<0.01 for uninfected versus infected; #p<0.05 for WT vs. *ATF4* KO by means of two-way ANOVA analysis. (**B**-**H**) ES-2 cells used for all experiments.

We noted that infection increased the expression of promoters of mitochondrial biogenesis including nuclear factor erythroid 2-related factor 2 (NRF2) and peroxisome proliferator-activated receptor-γ coactivator 1a (PGC1a) (Supp. Fig. 3A-B) (*14*). To therefore determine whether the increase in mtDNA was due to infection-induced mitochondrial biogenesis, we compared nuclear and mitochondrially encoded transcripts levels including *SDHB* and mt-ND1 in uninfected and *Toxoplasma-*infected cells. However, infection did not increase their levels (Supp. Fig. 3C-D). Consistent with these results, flow cytometry-based analysis of mitochondrial mass revealed minimal differences between uninfected and infected cells at 6, 12, and 24 hpi (Supp. Fig. 3E). Thus, mtDNA copy number is increased during infection independently of global changes in mitochondrial abundance.

How do *Toxoplasma-*infected cells increase mtDNA levels? We noted that infection also induced the expression of the mtDNA transcription factor *TFAM* and mtDNA helicase *TWNK,* whose mild overexpression increases mtDNA levels by ∼1.5x-fold, comparable to changes we observed during infection (Supp. Fig. 4, Fig. 1B) (*15*). We therefore posited that the increase in mtDNA was regulated by an infection-induced transcription factor upstream of *TFAM* and *TWNK*. To address this possibility, we compared the protein abundance in whole cell extracts from uninfected cells and *Toxoplasma-*infected cells using mass spectrometry (Supp. Table 1). Infection with *Toxoplasma* caused an increase in the abundance of several transcription factors including c-MYC, JUN, and FOS, in line with previous observations (Fig. 1C) (*16–18*). However, the activating transcription factor 4 (ATF4) was the highest induced host transcription factor at a ∼16x-fold enrichment in proteomes of infected cells (Fig. 1C).

ATF4 is the main effector of the integrated stress response (ISR), a common adaptive pathway that promotes cellular recovery during various stresses including ER stress and viral infection (*19*). It was of great interest to us because in recent years, ATF4 has been recognized as a key regulator of mitochondrial stress responses (*20*). Furthermore, the ISR promotes mtDNA recovery in cells with mtDNA double-stranded breaks (*21*). In confirmation of a link between the ISR and mtDNA regulation, we found that the treatment of cells with tunicamycin, an inhibitor of N-linked glycan biosynthesis that activates the ISR and thus ATF4, was sufficient to increase mtDNA levels independently of infection (Supp. Fig. 5). We therefore sought to address the role of ATF4 in mtDNA dynamics during infection. To first confirm our proteomics results, we analyzed uninfected cells and *Toxoplasma-*infected cells by immunoblot analysis. Infection induced ATF4 as early as 6 hpi and maximally at 24 hpi (Fig. 1D). To test the role of ATF4 in increasing mtDNA levels during infection, we generated ATF4 knockout (KO) ES-2 cells using CRISPR-Cas9 technology (Fig. 1E). The loss of ATF4 was sufficient to prevent the increase in mtDNA copy number during both *Toxoplasma* infection or tunicamycin treatment in wild-type (WT) cells (Fig. 1F-H; Supp. Fig. 5). Thus, ATF4 activation regulates mtDNA levels during infection and infection-independent stress.

### ATF4 increases mtDNA levels in a mitochondrial 1C metabolism-dependent manner

How does ATF4 regulate mtDNA during infection? The mild overexpression of *TFAM* and *TWNK* are sufficient to increase mtDNA to levels similar to those we observed during infection (*15, 22*). Because infection induced the expression of *TFAM* and *TWNK,* we posited that ATF4 regulated mtDNA levels in a TFAM- and TWNK-dependent manner. To address this possibility, we examined their expression in uninfected and *Toxoplasma-*infected WT and ATF4 KO cells.

Contrary to our expectation, the loss of ATF4 did not block the increase in *TFAM* and *TWNK* transcripts during infection (Supp. Fig. 4A-B). In fact, ATF4 KO cells exhibited higher expression of *TWNK* relative to WT uninfected and *Toxoplasma-*infected cells (Supp. Fig. 4A). These results indicated that ATF4 regulated mtDNA independently of TFAM and Twinkle, and raised the question of how ATF4 drove an increase in mtDNA during infection.

Because gene targets of ATF4 vary depending on its mechanism of activation, we next sought to define the host pathway upstream of ATF4 during infection (*23*). During the integrated stress response (ISR), diverse cellular stresses converge on the phosphorylation of eIF2α at serine 51, and the consequent attenuation of cap-dependent translation leads to the translation of ISR effectors such as ATF4 (*19*). Alternatively, ATF4 can be activated by pro-growth signals that stimulate mTORC1, the anabolic regulator of the cell (*24*). To determine whether the ISR or mTORC1 mediated ATF4 activation during infection, we generated mouse embryonic stem cells cells (mESCs) deficient for eIF2α phosphorylation (eIF2α^S51A^). In WT mESCs, both *Toxoplasma* infection and tunicamycin treatment drove eIF2α phosphorylation, and ATF4 induction as in ES-2 cells (Fig. 2A, Supp. Fig. 6A). However, neither induced ATF4 in eIF2α^S51A^ mESCs or in ES-2 cells treated with ISRIB, a small molecule inhibitor of the ISR (Fig. 2A; Supp. Fig. 6A). Consistent with this result, rapamycin- or torin1-based inhibition of mTORC1, as assessed by phosphorylation of its target ribosomal protein S6 kinase 1 (S6K1), did not block ATF4 activation during *Toxoplasma* infection (Supp. Fig. 6B). Thus, infection activates ATF4 in an ISR-dependent manner.

**Fig. 2.**
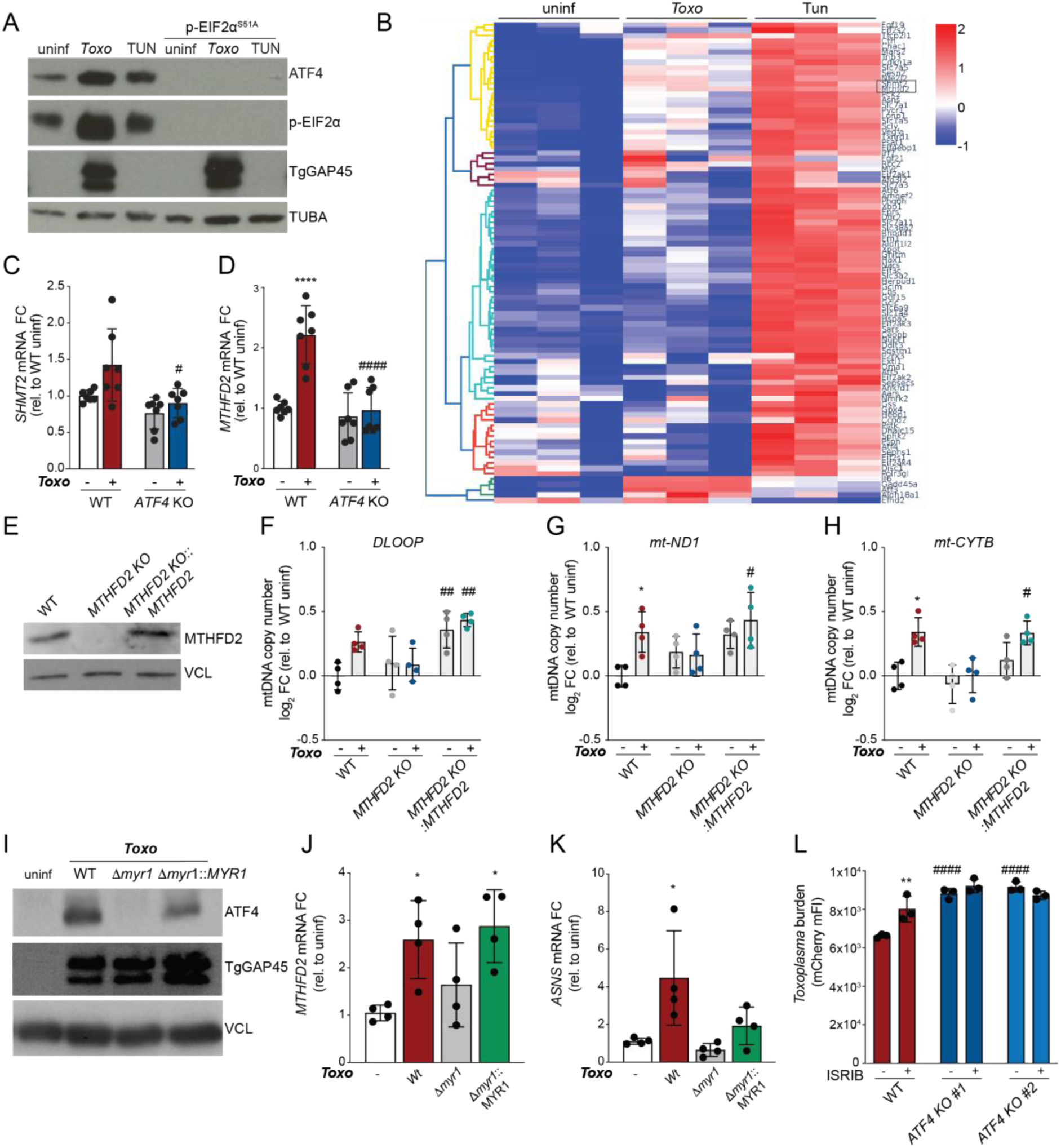
Host detection of parasite effectors leads to ISR-mediated ATF4 activation and an increase in mtDNA levels in a mitochondrial one-carbon metabolism-dependent manner. (**A**) Immunoblot (IB) analyses of lysates from WT and eIF2α^S51A^ AN3-12 mouse embryonic stem cells that were either uninfected, infected with *Toxoplasma*, or tunicamycin-treated (3 μg/ml) for 24 hours: ATF4, ∼50 kDa; phospho-EIF2α (Ser51), ∼38 kDa; α-Tubulin (TUBA), ∼55 kDa; and *Toxoplasma* GAP45 (TgGAP45), ∼45 kDa. (**B**) Heatmap of mRNA levels of *ATF4* target genes in uninfected, infected and tunicamycin (3 μg/ml) treated ES-2 cells; n=3 independent cultures; *SHMT2* and *MTHFD2* are boxed. (**C** and **D**) WT and *ATF4* KO ES-2 cells were uninfected or infected with *Toxoplasma* for 24 h and analyzed by qPCR for *SHMT2* and *MTHFD2*. Transcript levels were normalized to *ACTB* and are relative to WT uninfected. Data are mean ± SD of n=7 independent cultures. ****p<0.0001 for uninfected versus infected by means of two-way ANOVA analysis. (**E**) IB of ES-2 cells of the indicated genotype: MTHFD2, ∼32 kDa; α-Tubulin (TUBA), ∼55 kDa. (**F, G** and **H**) ES-2 cells of the indicated genotype were analyzed for mtDNA by qPCR for *DLOOP*, mt*-ND1* and mt*-CTYB* normalized to *RUNX2* levels and relative to WT uninfected. Data are mean ± SD of n=4 independent cultures. *p<0.05; for uninfected versus *Toxoplasma* infected, and #p<0.05; ##p<0.01 for WT versus *MTHFD2* KO by means of two-way ANOVA analysis. (**I**) IB analysis of lysates from ES-2 cells that were uninfected or infected with indicated *Toxo* strains (MOI: 4) at 24 hpi: ATF4, ∼50 kDa; Vinculin (VCL), ∼124 kDa; and *Toxoplasma* GAP45 (TgGAP45), ∼45 kDa. (**J** and **K)** Cells treated as in (**I**) were analyzed by qPCR for *MTHFD2* and *ASNS*. Transcript levels were normalized to *ACTB* levels and are relative to uninfected cells. Data are mean ± SD of n=4 independent cultures, *p<0.05; **p<0.01 by means of one-way ANOVA analysis. (**L**) WT and *ATF4* KO ES-2 were infected with *Toxoplasma* ± ISRIB (200nM) and analyzed 24 hpi by means of flow cytometry for *Toxoplasma* burden (mCherry median FI). Data are mean ± SEM of n=3 biological experiments, **p< 0.01 for DMSO versus ISRIB, ####*p* < 0.0001 for WT versus ATF4 KO by means of two-way ANOVA analysis. (**B**-**L**) ES-2 cells used for all experiments.

To next identify the ATF4 targets common to infection and ISR activation that regulated mtDNA dynamics, we compared the expression of a panel of ATF4 targets in uninfected cells, *Toxoplasma-*infected cells, and cells treated with tunicamycin (Supp. Table S2) (*25*). ATF4 targets induced by both *Toxoplasma* infection and tunicamycin treatment included the transcription factor *ATF3* and asparagine synthase (*ASNS*) (Fig. 2B). Of particular interest were *MTHFD2* and *SHMT2*, enzymes of the mitochondrial arm of the 1C metabolism processes that use folate as cofactor to activate and transfer 1C groups for the biosynthesis of nucleotide precursors (Fig. 2B) (*7, 20, 25, 26*). Furthermore, MTFHD2 has been linked to mtDNA deficiencies (*8, 26*). As expected, we found that infection induced *MTHFD2* and *SHMT2* in an ATF4-dependent manner (Fig. 2C-D). To test the importance of mitochondrial 1C (mito-1C) metabolism in the changes in mtDNA during infection, we generated cells deficient for MTHFD2 (Fig. 2E). Infection did not increase mtDNA levels in MTHFD2 KO cells, unlike in WT cells (Fig. 2F-H). Of note, the overexpression of *MTHFD2* cDNA in MTHFD2 KO cells rescued the increase in mtDNA levels in infected cells, and was sufficient to increase mtDNA levels in uninfected cells (Fig. 2F-H). Thus, mito-1C metabolism is required for ATF4 to increase mtDNA levels during infection.

### Host cells activate ATF4 in response to the detection of secreted parasite effectors

Because ATF4 regulated mtDNA dynamics in a mito-1C metabolism-dependent manner, we hypothesized that the activation of ATF4 comprised a defensive host response to *Toxoplasma.* However, the possibility remained that the ISR was activated by metabolic stress caused by the siphoning of nutrients by replicating parasites, as previously suggested (*16*). To distinguish between these possibilities, we asked if the induction of ATF4 required parasite effector proteins that are secreted into the host cell. To do so, we examined ATF4 activation in cells infected with Δ*myr1* parasites that lack the translocon apparatus required for the secretion of a class of effector proteins, but are competent for replication and thus nutrient siphoning (*27, 28*). Host cells infected with Δ*myr1* parasites did not induce ATF4 nor its targets *MTHFD2* and *ASNS* (Fig. 2I-K). Consistent with our hypothesis that ATF4 activation comprised a host defense, both the pharmacological suppression of ATF4 via ISRIB and the genetic ablation of ATF4 led to significant increases in parasite replication (Fig. 2L; Supp. Fig. 6A). Thus, the activation of ATF4 in response to host detection of *Toxoplasma* effectors restricts parasite growth.

### ATF4 drives mitochondrial 1C-metabolism to limit parasite dTMP synthesis and proliferation

How does ATF4 induction affect mitochondrial metabolism and parasite growth? To address this question, we first assessed the contribution of mito-1C metabolism to dTMP and dTTP biosynthesis during *Toxoplasma* infection. Because serine is the predominant donor of carbons to 1C pathways in cultured cells, we cultured uninfected and *Toxoplasma-*infected WT cells with deuterium-labelled serine [2,3,3-^2^H-serine], and examined the resulting incorporation of ^2^H into dTMP or dTTP (Fig. 3A) (*7, 29*). The contribution of mito-1C-derived folates leads to dTMP and dTTP with 1 deuterium and a mass shift of 1Da (M+1); the contribution of cytosolic-1C (cyto-1C) -derived folates will instead lead to the incorporation of 2 deuterium atoms and results in M+2 dTMP or dTTP (Fig. 3A) (*29*). In uninfected WT cells, M+1 dTMP and M+1 dTTP were the dominant isotopologes at 100% and >98% of their respective isotope-labeled pools (Fig. 3B-C). This result is in line with previous observations that mito-1C predominately supplies 1C units in most growing cell lines (*29*). However, *Toxoplasma* infection of WT cells led to a 10x-fold increase in M+2 dTTP levels; M+2 dTMP was not detected, likely due to the lower abundance of intracellular dTMP pools (Fig. 3B-C). We next assessed the effect of ATF4 ablation on mito-1C metabolism during infection. Remarkably, we found that the loss of ATF4 led to a significant increase in the contribution of cyto-1C to dTMP and dTTP synthesis. M+2 dTMP, which was not detected in WT infected cells nor uninfected ATF4 KO cells, made up ∼20% of labelled dTMP in infected ATF4 KO cells, while M+2 dTTP was ∼2x-fold higher than infected WT cells (Fig. 3B-C). Thus, ATF4 maintains host mito-1C metabolism during infection.

**Fig. 3.**
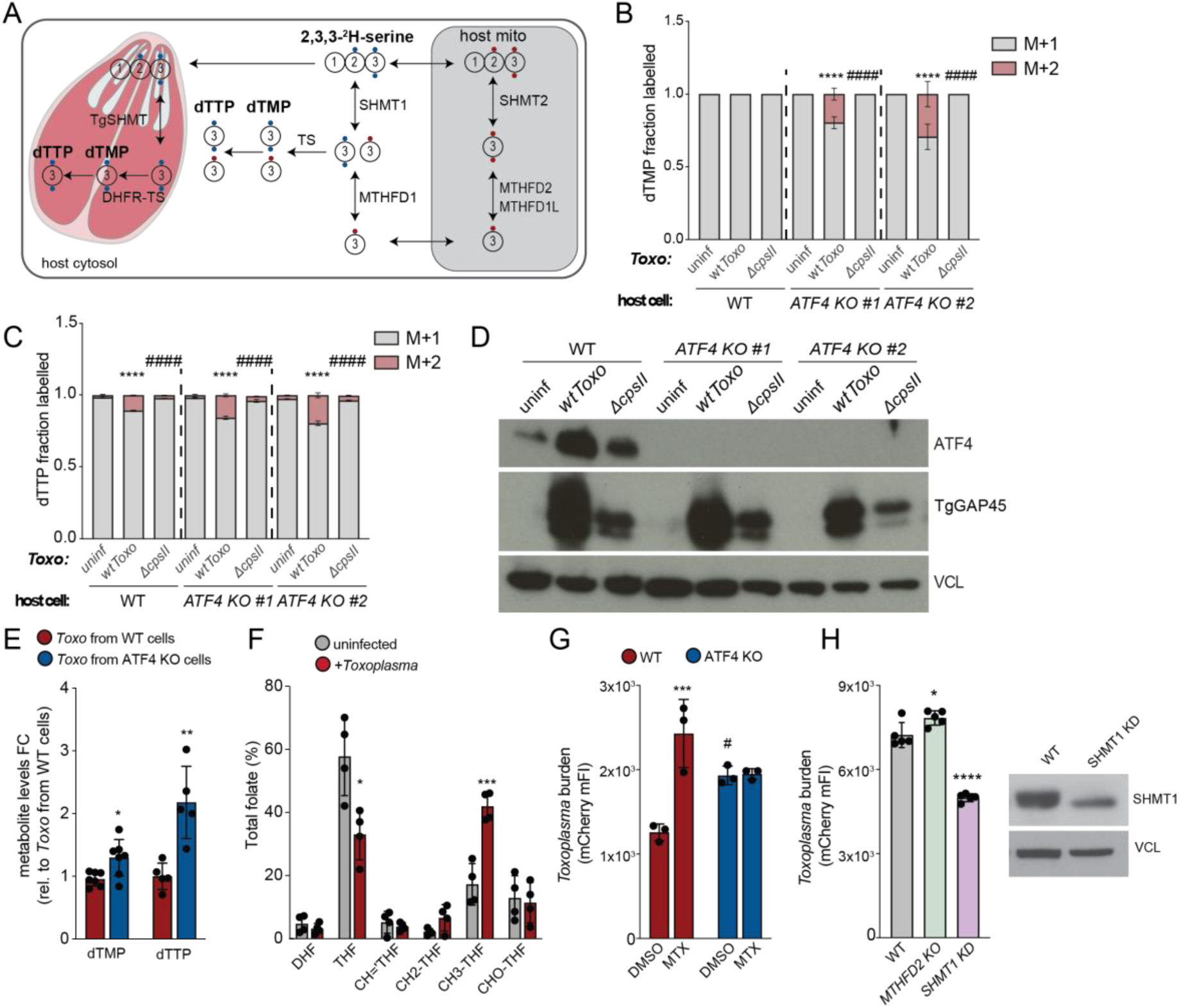
ATF4 drives host use of folate to limit parasite dTMP synthesis and parasite replication. (**A**) Schematic of parasite and host one-carbon metabolism as determined by incorporation of ^2^H from [2,3,3-^2^H] serine-labelling into dTMP and dTTP. Contribution of cytosolic-1C (cyto-1C) metabolism to dTMP production results in M+2 ^2^H-labelleled dTMP (2 blue dots), whereas its production from 1C units generated in mitochondria produces M+1 ^2^H-labelled dTMP (1 red dot). *Toxoplasma* has a cytosolic SHMT and thus parasite-1C metabolism generates M+2 ^2^H-labelleled dTMP. (**B** and **C**) dTMP and dTTP labeling for 24h during infection in WT and two clones of *ATF4 KO* ES-2 cells uninfected or infected with WT *Toxoplasma* and Δ*cpsII* (Carbamoyl phosphate synthase II) parasites. Data are mean ± SD of n=3 independent cultures, ****p<0.0001 for uninfected versus infected and ####p<0.0001 for WT versus ATF KO clones by means of two-way ANOVA analysis. (**D**) ES-2 cells with indicated genotypes were uninfected or infected with WT *Toxoplasma* or Δ*cpsII* parasites and harvested at 24 hpi for analysis by immunoblotting for ATF4, ∼50 kDa; Vinculin (VCL), ∼124 kDa; and *Toxoplasma* Gap45 (TgGAP45), ∼45 kDa. (**E**) Abundance of *Toxoplasma* dTMP levels in WT-*Toxoplasma* infected cells and *ATF4* KO-infected cells for 24hpi. Data are mean ± SD of n=7 independent cultures, dTTP not detected in two cultures, *p<0.05; **p<0.01; for WT versus *ATF4* KO by means of unpaired t-test. (**F**) Total levels of folate represented as (%) were measured by UPLC-MS/MS in uninfected and *Toxoplasma*-infected ES-2 cells at MOI=4 at 24 hpi and normalized to cell number. DHF: dihydrofolate; THF: tetrahydrofolate; CH=‘THF: 5,10-methenyl-tetrahydrofolate; CH_2_-THF: 5,10-methylene-tetrahydrofolate; CH_3_-THF: 5-methyl-tetrahydrofolate; CHO-THF: 10-formyl-tetrahydrofolate. Data are ± SD of n=4 independent cultures *p<0.05; ***p<0.001 for uninfected versus infected by means of multiple unpaired t-test. (**G**) WT and ATF4 *KO* ES-2 cells were infected with *Toxoplasma* ± MTX (200 nM) and analyzed 24 hpi by means of flow cytometry for *Toxoplasma* burden (mCherry median FI). Data are mean ± SEM of three biological experiments, ***p < 0.001 for DMSO versus MTX, and #*p* < 0.05 for WT versus ATF4 *KO* by means of two-way ANOVA analysis. (**H**) WT, MTHFD2 KO, and SHMT1 knockdown (KD) ES-2 cells were infected with *Toxoplasma* and analyzed as in (**G**). Data are mean ± SEM of five biological experiments, *p<0.05; ****p< 0.001; by means of one-way ANOVA; IB analyses of lysates from WT and *SHMT1* KD cells: SHMT1, ∼55 kDa; VCL, ∼124 kDa. (**B-H**) ES-2 cells used for all experiments.

We reasoned that the observed increase in M+2 dTMP and dTTP in infected WT and ATF4 KO cells could result from an increased reliance on host cyto-1C. Alternatively, parasite-derived dTMP and dTTP synthesis could contribute to the increase in M+2 dTMP and dTTP isotopologues, given the fact that *Toxoplasma* expresses a single cytosolic SHMT and can synthesize pyrimidines de novo (Fig. 3A) (*30, 31*). To distinguish between these possibilities, we first assessed levels of pyrimidines in uninfected and *Toxoplasma-*infected cells treated with or without leflunomide, a small molecule inhibitor of host dihydroorotate dehydrogenase and thus pyrimidine synthesis (*32*). We found that infection led to ∼10x-fold and ∼2x-fold increases in total dTMP and dTTP, even in the presence of leflunomide, despite the expected accumulation of the pyrimidine precursors carbamoyl aspartic acid and dihydroorotic acid (Supp. Fig. 7A-B). This result indicated that the increases observed during infection were, in part driven by *Toxoplasma*-derived dTMP and dTTP.

We next addressed whether the increase in M+2 dTMP and dTTP in ATF4 KO cells was due to their synthesis by host cells or *Toxoplasma*. To do so, we examined M+1 and M+2 isotopologues of dTMP and dTTP during infection with parasites that are deficient for their synthesis de novo due to the loss of carbamoyl phosphate synthetase II (CPSII), and thus only replicate if supplemented with exogenous uracil (*30*). Infection with *ΔcpsII* parasites led to ATF4 activation, but unlike in the case of WT parasites, failed to drive the increases in M+2 dTMP and M+2 dTTP observed in ATF4 KO cells (Fig. 3B-D). These results indicated that the M+2 dTMP and M+2 dTTP detected in infected ATF4 KO cells were parasite-derived, and that the loss of ATF4 promoted parasite dTMP synthesis. In line with this result, parasites isolated from ATF4 KO cells had higher total dTMP and dTTP levels than parasites isolated from WT cells, but similar levels of host-derived dTMP (Fig. 3E; Supp. Fig. 8). Thus, ATF4 restricts parasite synthesis of dTMP.

How does ATF4 activation regulate parasite dTMP levels? Unlike in mammals, in *Toxoplasma,* the synthesis of dTMP is catalyzed by the bifunctional enzyme dihydrofolate reductase thymidylate synthase (DHFR-TS) (Supp. Fig. 7C) (*33*). This means that parasites couple the generation of dTMP, the rate-limiting step in DNA synthesis, to the reduction of dihydrofolate (DHF) to tetrahydrofolate (THF). To catalyze these reactions, *Toxoplasma* therefore requires both DHF, the precursor of all folates, and its derivative 5, 10 methylene-THF (5,10-CH2-THF) (Supp. Fig. 7C). Because ATF4 activation induced the mito-1C cycle that uses folates, we reasoned that ATF4 promoted mitochondrial use of folates, thereby restricting *Toxoplasma* access to the folates required for dTMP synthesis and parasite growth. To address this possibility, we assessed levels of folates in uninfected and infected cells using mass spectrometry. Infection led to a ∼50% decrease in THF, and a slight decrease in its precursor DHF, showing that flux through the folate cycle is altered during infection which, may limit availability of certain folates (Fig. 3F). To next test whether ATF4 restricted parasite growth in a folate-dependent manner, we treated WT and ATF4 KO cells with methotrexate (MTX), an inhibitor of host dihydrofolate reductase (DHFR) that converts dietary folic acid into DHF and THF for use in 1C metabolism reactions. Of note, MTX is not taken up by *Toxoplasma,* and thus does not inhibit *Toxoplasma* DHFR-TS (*34*). We found that MTX treatment increased parasite proliferation in WT cells, but not in ATF4 KO cells (Fig. 3G). Furthermore, impairing mito-1C metabolism through ablation of the ATF4 target MTHFD2 or knockdown of the reduced folate carrier SLC19A1 rendered cells permissive to parasite replication and increased dTMP levels (Fig. 3H, Supp. Fig. 9) (*35*). Conversely, driving host-dependence on mito-1C metabolism by genetic ablation of the cyto-1C enzyme SHMT1 restricted parasite growth (Fig. 3H). These results indicate that ATF4 promotes host mitochondrial use of folates, thereby restricting parasite dTMP synthesis and growth.

### ATF4 activates a host-protective response in vivo

Our findings that infected cells activate mito-1C metabolism to restrict parasite growth led us to ask whether ATF4 activated a host-protective response in vivo. To address this question, we first asked whether ATF4 was activated in a mouse model of *Toxoplasma* infection. We therefore injected mice intraperitoneally with vehicle, tunicamycin, or mCherry-expressing *Toxoplasma* and measured the levels of ATF4 target transcripts in peritoneal exudate cells (PECs) isolated near the peak of acute infection (5 dpi) (*36*). Infection drove increases in the levels of ATF4 targets *Atf3, Shmt2*, and *Mthfd2*, similar to our results in cultured cells (Fig. 2B; Supp. Fig. 10). To determine whether the observed increases during infection were due to ISR-mediated activation of ATF4, mice were injected with vehicle or the ISR inhibitor ISRIB prior to infection and after every other day (Fig. 4A). ISRIB treatment blunted the induction of ATF4 targets *Atf3, Mthfd2* and *Shmt2* levels observed in PECs isolated from infected mice (Fig. 4B-D). We next tested whether blocking ATF4 activation affected *Toxoplasma* proliferation in vivo. To this end, we examined the *Toxoplasma* burden in peritoneal exudate of ISRIB-versus vehicle-treated mice. We found that ISRIB treatment led to higher total *Toxoplasma* levels as assessed by qPCR, which was due to an increase in extracellular parasites (Fig. 4E-H). Taken together, these data indicate that ATF4 restricts parasite proliferation in vivo.

**Fig. 4.**
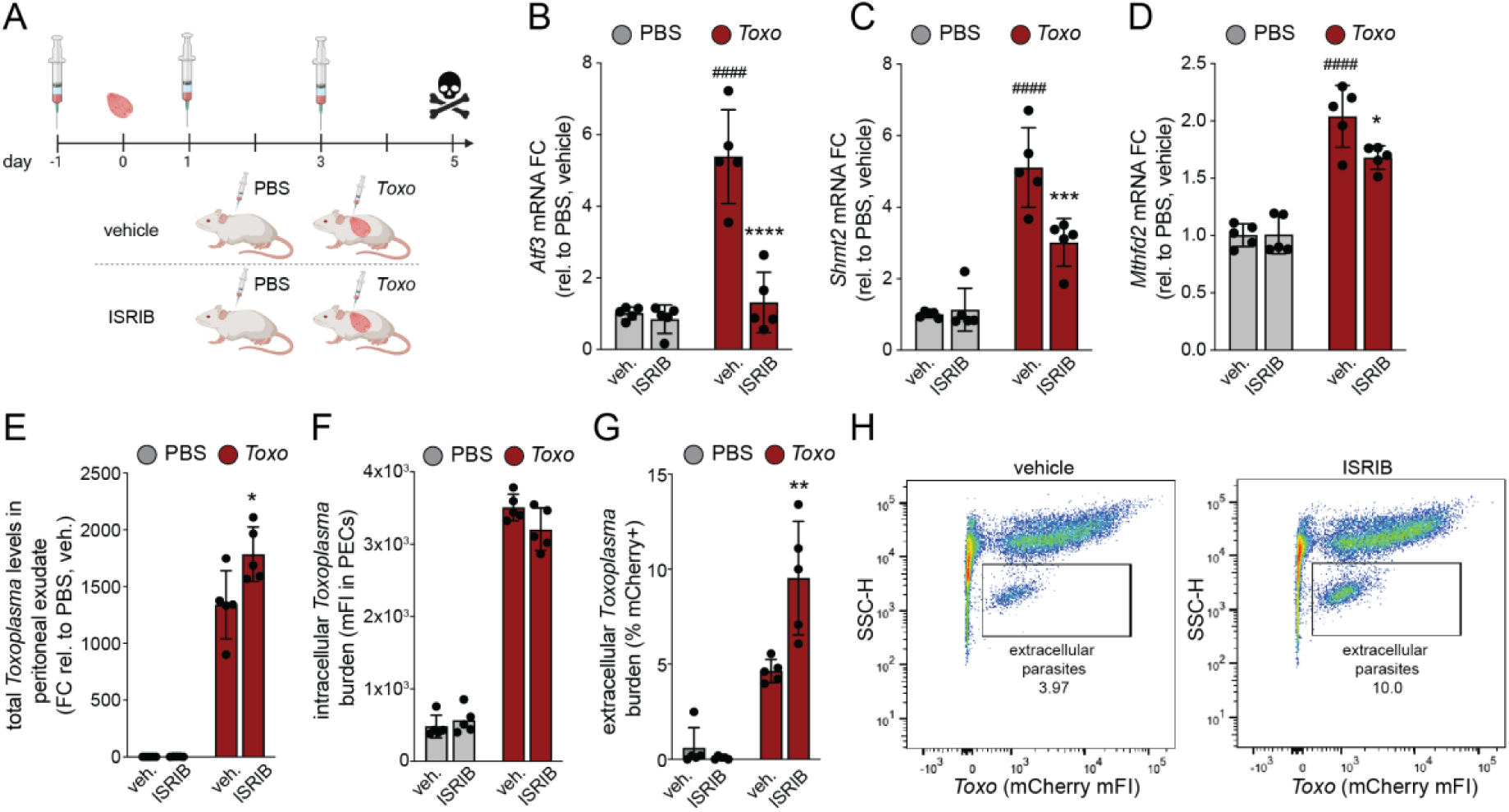
ATF4 activates a host-protective response in vivo. (**A**) Schematic of infection and ISRIB administration. Mice (n=5) where injected with ISRIB one day before infection with 50 tachyzoites or 1x PBS injection and every other day after (**B-D**) Peritoneal exudate cells (PECs) were isolated from mice treated as in (**A**) at 5 days post injection (dpi) and analyzed for the indicated ATF4 targets genes by qPCR. Transcripts are normalized to *Hprt* and relative to uninfected PECs mice (n=5). *p<0.05; ***p<0.001; ****p<0.0001 for vehicle versus ISRIB and #p < 0.05; ###p < 0.001; and ###p < 0.0001 for PBS versus *Toxoplasma* by means of two-way ANOVA analysis. (**E**) Peritoneal exudate was isolated 5 dpi from mice treated as in (**A**) and analyzed by qPCR for *Toxoplasma SAG1* levels. DNA levels were normalized to *Hprt.* Data are mean ± SD of n=5 mice, *p<0.05 by unpaired t-tests. Peritoneal exudate isolated from mice treated as in (**A**) was analyzed by means of flow cytometry for (**F**) Intracellular *Toxoplasma* parasite burden (mCherry median FI) and (**G**) percentage of extracellular parasites. Data are mean ± SD of n=5 mice, *p<0.05; **p<0.01 by unpaired t-test analysis (vehicle vs. ISRIB). (**H**) Representative scatter plots of peritoneal exudate isolated from mice treated as in (**A**).

Our data shows that ATF4 effects a folate-based metabolic defense against *Toxoplasma* that involves increased activity of mitochondrial 1C metabolism. This raises several questions, beginning with how do host cells activate ATF4? The secretion of effector proteins was required for ATF4 induction via the ISR. A *Toxoplasma* effector(s) could therefore directly be sensed by one of the kinases that phosphorylate eIF2α, or cause damage that indirectly triggers ISR activation.

1C metabolism processes occur in the cytosol and mitochondria. So why is mito-1C metabolism, but not cyto-1C metabolism host-protective? Although *Toxoplasma* is capable of folate synthesis, it also scavenges exogenous folate, suggesting this metabolite is required in significant abundance for parasite proliferation (*37*). Unlike in mammals, in *Toxoplasma* folate availability is tied to the production of dTTP, the rate-limiting step in DNA synthesis that is required for the proliferation of any organism (Supp. Fig. 7C). *Toxoplasma* however appears deficient for thymidine salvage (*11*). It is therefore likely that the parasite is completely dependent on DHFR-TS for the dTTP required for its DNA replication. Compartmentalization of 1C metabolism processes within the mitochondria rather than the cytosol would block parasite access to folates it requires for growth. In line with this expectation, the loss of ATF4 and thus the corresponding changes in mito-1C metabolism led to an increase in parasite dTMP synthesis and parasite burden (Fig. 2L, 3B-E). Furthermore, impairing host use of folate for 1C-metabolism processes via methotrexate (MTX), a drug that is widely used in anti-cancer therapies, led to increased growth in WT cells but not ATF4 KO cells (Fig. 3G). These results support a role for mito-1C metabolism in limiting parasite access to folate.

The question also arises of whether promoting host mitochondrial use of folate may suppress the growth of *Toxoplasma* in vivo, or other human pathogens that require folate for dTMP synthesis such as *Cryptosporidium hominis, Leishmania major,* and *Plasmodium falciparum,* the causative agent of malaria. Conversely, does increasing folate availability through impairing host use or dietary supplementation without a parallel modulation of mito-1C metabolism enhance replication of these pathogens? MTX administration during acute *Toxoplasma* infection converted nonlethal infections into severe disease; whether increased parasite proliferation was a contributing factor to the worsening of disease is unknown (*38*). High dietary folate increased parasite replication in a murine malaria model, and increased folate concentrations were associated with an increase in malaria infection in humans (*39, 40*). These observations hint at a key role for folate competition in certain host-pathogen interactions.

Last, does mito-1C metabolism enable mitochondria to compete for metabolites beyond folate? Mitochondria and intracellular pathogens both depend on cytosolic nucleotides for the replication of their DNA, and thus ostensibly compete for the same dNTP pools. The nucleotide burden to replicate one 16 kB molecule of mtDNA seems negligible. However, given that each of the estimated ∼880 mitochondria per cell contain on average 3 genomes, the total mtDNA burden per cell instead is ∼42 MB (*1*). Enhancing mito-1C metabolism may promote their synthesis proximal to mitochondria, and thus facilitate mitochondrial uptake of nucleotides. Although we focused here on the human pathogen *Toxoplasma,* which is a protozoan parasite, our findings parallel those from studies with diverse pathogens that rely on host nucleotides but not host folate. For example, the ablation of the cyto-1C enzyme SHMT1 reduced proliferation of SARS-CoV-2, the causative agent of COVID-19, while the loss of the analogous mito-1C enzyme SHMT2 had the opposite effect (*41*).

Our study of the interaction between the human parasite *Toxoplasma* and host mitochondria led to the discovery of ATF4 as an effector of a folate-based metabolic immunity that enables host cells to restrict parasite growth, rather than simply functioning as a stress sensor. Furthermore, our findings demonstrate the importance of mitochondrial metabolism in protecting host cells during infection and shed light on cell autonomous defenses against pathogens. This paves the way for future studies exploring organelle-mediated defense strategies and the role of folate competition during infectious diseases.

## Acknowledgments

We thank E. Cors and T. Langer (MPI-AGE) for the generous sharing of reagents; the MPI-AGE Metabolomics Core, in particular F. Dethloff, C. Edlich-Muth and Y. Hinze for advice on sample preparation and Shiny App support for metabolite analysis; the MPI-Proteomics facility; the MPI-AGE FACS and Imaging Core for flow cytometry and microscopy support; the MPI-AGE Comparative Biology Core Facility, in particular S. Buhl and B. Bertalan. We also thank P. Krueger, J. Straub and A. Lamprakis for manuscript feedback, and all members of the Pernas laboratory for helpful discussions.

## Author contributions

Conceptualization: T.C.M and L.F.P. Methodology: T.C.M., S.R., X.L., B.G.M., I.R., K.L., N.G., M.P., A.M., P.G., and L.F.P. Investigation: All authors. Resources: K.A., M.D.H., M.S.D., Nanostring analysis: T.C.M, Proteomics analysis: L.F.P., Funding acquisition: L.F.P. Project administration: L.F.P. Writing, original draft: T.C.M. and L.F.P. Review and editing: All authors. Supervision: L.F.P.

## Data and materials availability

All data are available in the main text or the supplementary materials.

## List of Supplementary Materials

Materials and Methods

Supplementary Figures 1-10

Supplementary Table 1-3 (Proteomics; Nanostring Data, List of primers)

## Supplementary Materials

**Other Supplementary Materials for this manuscript include the following:**

Supplementary Tables S1-S3

### Materials and Methods

#### Cell culture and cell lines

HeLa adenocarcinoma cells, ES-2 ovary clear cell carcinoma, Human foreskin fibroblasts (HFF), were obtained from ATCC (CCL-2, CRL-1978, and SCRC-1041, respectively); ES-2 cells stably expressing pMXs-eGFP-OMP25 (referred to as OMM-GFP; Addgene #83356) were generated through lentiviral transduction. Unless stated otherwise cell lines were cultured in Dulbecco’s Modified Eagle’s Medium (DMEM) containing 10% heat-inactivated fetal bovine serum (FBS) and maintained at 37°C and 5% CO_2_. AN3-12 mouse embryonic stem cells were obtained from Dr. Josef Penninger (42) and were cultured in DMEM high glucose supplemented with glutamine, 15% fetal bovine serum, penicillin/streptomycin, nonessential amino acids, sodium pyruvate, β-mercaptoethanol, and LIF as previously described (43). ATF4 KO ES-2s were cultured in DMEM high glucose, 10% fetal bovine serum, β-mercaptoethanol and nonessential amino acids. Cells were tested every 2 weeks for Mycoplasma infection by means of polymerase chain reaction (PCR).

#### Parasite culture and strains

Toxoplasma gondii parasites of the Type I RHΔku80:mCherry+ (previously described in (44)); Type I RHΔmyr1 parasites and RHΔmyr1::Myr1 were provided by Dr. Moritz Treeck, (Gulbenkian Institute) (45); Type I RHΔcpsII:mCherry parasites were provided by Dr. David Bzik (Dartmouth Geisel School of Medicine) (46). Parasites were maintained by serial passage in human foreskin fibroblast (HFF) monolayers in cDMEM. RHΔcpsII: mCherry were maintained by serial passage in human foreskin fibroblast (HFF) monolayers in cDMEM supplemented with 0.2 mM uracil.

#### CRISPR/Cas9-mediated gene editing

Guide sequences used can be found in Supp. Table 3. ATF4 and MTHFD2 knockout (KO) cells and SHMT1 and SLC19A1 knockdown (KD) cells were generated using CRISPR-Cas9 mediated gene editing via lentiviral transductions into the pLenti CRISPRv2 (Addgene #5296). For production of lentiviral particles, 293T human embryonic kidney (HEK) cells were transduced using the X-tremeGENE 9 DNA Transfection Reagent (Roche) with 1 μg psPAX2 packaging vector (Addgene #12260), 0.3 μg pCMV-VSVG envelope vector (Addgene #8454) and 1 μg of the relevant plasmid of interest. The next day, the medium was exchanged, and two days post transfection, the virus-containing supernatant was filtered with a 0.45 μM filter and supplemented with polybrene to a final concentration of 5 μg/ml. Virus-containing medium was added to the indicated cell lines for 24h and puromycin selection (3 μg/ml) was started after an additional 24–48 h. Polyclonal cultures and individual clones were validated by immunoblotting or qPCR.

The eIF2α S51A substitution was engineered in AN3-12 cells using CRISPR/Cas9 technology as described previously (47). DNA template sequences for small guide RNAs were designed online (http://crispor.org), and cloned into the Cas9-mCherry expressing plasmid (Addgene #21852). Corresponding guide and Cas9 expressing plasmids were co-transfected with a single-stranded DNA repair template (Integrated DNA Technologies), using Lipofectamine 2000 (Thermo Fisher Scientific) according to the manufacturer’s instructions. mCherry-positive cells were singled in 96-well plates using FACSAria Fusion sorter and subjected to genotyping. DNA was extracted (DNA extraction solution, Epicentre Biotechnologies) and edited regions were specifically amplified by PCR. The PCR product was subjected to a restriction digest to identify positive clones. These clones have a gain of BglII restriction site due to a mutation in the repair template. Sanger sequencing was performed at Eurofins Genomics GmbH (Ebersberg, Germany). Positive clones were sorted prior to further experiments.

#### MTHFD2 overexpression

Human MTHFD2 cDNA amplified from ES-2 cells was cloned into pMSCV PIG (puro IRES GFP; Addgene #21654) at the XhoI and EcoRI sites. Following production of viral particles and transduction, puromycin selected ES-2 cells were sorted for GFP expression using flow cytometry using BD FACSDiva software.

#### Flow cytometry analysis

To assess mitochondrial mass, monolayers of ES-2 cells expressing OMM-GFP were rinsed with PBS, trypsinized and fixed in 2% paraformaldehyde in FACS buffer (3% FBS in 1XPBS) for 10 min. After a brief spin, cells were resuspended in FACS buffer and 10,000 events were analyzed on a FACSFortessa for GFP median fluorescence intensity (mFI) using BD FACSDiva software. To assess parasite proliferation, monolayers of ES-2 cells infected with RFP expression parasites (RHΔku80:mCherry+) were rinsed with PBS, trypsinized and fixed in 2% paraformaldehyde in FACS buffer (3% FBS in PBS) for 10 min. After a brief spin, cells were resuspended in FACS buffer and 10.000 events analyzed on a FACSFortessa and the RFP mean fluorescence intensity (mFI) using BD FACSDiva software.

#### Quantitative PCR for mtDNA analysis

For mtDNA measurements, genomic DNA was isolated from cell pellets that were washed once with 1x PBS using the Blood and Tissue DNA extraction kit (Qiagen #69504). DNA was quantified by nanodrop. 50 ng of genomic DNA were amplified using PowerSYBR Green PCR Master Mix (Thermo Fisher Scientific #A25742). Reactions were performed in 384 Hard-Shell microplates (BioRAD #HSP3801) sealed with adhesive qPCR seal (Roche) in a and Bio-Rad Real-Time PCR system with the following cycler program: 95°C for 7 min followed by 35 cycles of 95°C for 10 s, 60°C for 30 s. Dissociation curves confirmed single PCR products, and signals were analyzed within the linear amplification range. Each sample was run in triplicate. Relative mtDNA copy number changes were calculated using the comparative ΔΔCt method (48) by determining Ct threshold values and using equations ΔΔCt = (Ct_mtDNA − Ct_control) at t(n) − (Ct_mtDNA − Ct_control) at t(0) and 2^−ΔΔCt^. The primers DLOOP, mt-ND1, mt-CYTB, mt-ND6 and mt-COX1 as mitochondrial genes and RUNX2 and ACTB as nuclear DNA controls can be found in Supp. Table 3.

#### Quantitative PCR for mRNA analysis

For analysis of mRNA expression, RNA purification was performed using TriZol (Thermo Fischer Scientific; #15596018) according to the manufacturer’s instructions. The Reverse Transcriptase reaction (RT) was done using 1 μg of RNA for each sample into cDNA using SuperScript™ VILO™ Master Mix (Thermo Fischer Scientific; # 11755050). For each reaction, 2.5 ng of cDNA were amplified using PowerSYBR Green PCR Master Mix and Bio-Rad Real-Time PCR system. For each independent sample, RT–qPCR was performed in technical triplicates. The primer sequences used in this study can be found in Supp. Table 3. Expression levels were calculated by the delta delta ct methods as described above, for which ACTB and HPRT was used as a control. Relative fold changes were calculated using the comparative ΔΔCt method (48) by determining Ct threshold values and using equations ΔΔCt = (Ct_targetgene − Ct_control) at t(n) − (Ct_targetgene − Ct_control) at t(0) and 2^−ΔΔCt^.

#### Immunoblotting and antibodies

Whole cells were harvested in chilled lysis buffer (50mM Hepes-KOH pH 7.4, 40mM NaCl, 2mM EDTA, 1.5mM NaVO4, 50mM NaF, 10mM NaPyrophosphate, 10mM, NaBetaGlycerophosphate (disodium salt pentahydrate), 1% Triton X-100) and lysed for 30 min on ice. Lysates were subsequently centrifuged at 10 min at 14,000 x g at 4°C and the clarified supernatant was transferred into a fresh tube with 5X SDS for a final volume of 1X SDS. Following SDS-PAGE and gel transfer, membranes were blocked with TBS-0.05% Tween 20 (TBS-T) and 5% milk and the primary antibodies were incubated overnight. Following incubation, blots were washed three times in TBS-T and then incubated with horseradish peroxidase (HRP)-conjugated anti-mouse IgG (CST #7076) or anti-rabbit IgG (CST #7074) at a 1:4000 dilution for 45 minutes and developed using a chemiluminescence system (Pierce™ ECL Western Blotting Substrate or Pierce SuperSignal™ West Atto Ultimate Sensitivity Substrate; ThermoFisher Scientific). The following antibodies were used: ATF4 (CST #11815), VCL (CST #4650), pS6K (CST #97596), TUBA (Proteintech 66031-1-Ig), MTHFD2 (CST #41377), SHMT1 (CST #80715), MTHFD1L (Invitrogen #PA5-100158), MTHFD1 (Sigma #HPA015006), phospho-eIF2a (Ser51) (CST #9721), ACTB (CST #4970), TgGra45 (Dr. D Soldati; U. of Geneva) and TgGRA7 (Dr. J Boothroyd; Stanford University).

#### Live-cell imaging

Cells were plated on 6-well CELLview glass bottom cell culture dishes (Greiner Bio-One). The following morning, cells were infected with RH mcherry expressing Toxoplasma. After 24 hours, cells were incubated with MitoTracker Deep Red (Thermo Fisher Scientific, 50 nM) and Picogreen 1ug/ml for 30 min and imaged using an Olympus IXplore SpinSR 50 mm spinning disk confocal microscope. Live cell imaging was performed in cDMEM with incubation at 37°C and 5% CO_2_. All images were taken with a 100X/1.35 silicon oil objective and excitation with either 488, 561, or 640 laser lines, using ORCA-Flash4.0 cameras (Hamatsu), and cellSens Software.

#### Proteomics sample preparation

As previously described in (49), for preparing samples from immunoprecipitation, on-beads digestion was performed to elute the proteins off the beads. Before adding the elution buffer, the beads were washed with detergent-free buffer (50mM Tris-HCl pH7.5) four times to remove any detergents used previously. Then 100 µl of the elution buffer (5ng/µl trypsin, 50mM Tris-HCl pH7.5, 1mM Tris(2-carboxyethyl)phosphine), 5mM chloroacetamide) was added to the beads and incubated at room temperature by vortexing from time to time, or rotating on a rotator. After 30 min, the supernatant was transferred to a 0.5 ml tube and incubated at 37°C overnight to ensure a complete tryptic digest. The digestion was stopped in the next morning by adding formic acid to the final concentration of 1%. The resulted peptides were cleaned with home-made StageTips. Alternatively, four micrograms of the eluted peptides were dried out and reconstituted in 9 µL of 0.1M TEAB and labeled with tandem mass tags (TMTpro, Thermo Fisher Scientific cat. No A44522). Labeling was carried out according to manufacturer’s instruction with the following changes: 0.5 mg of TMTPro reagent was re-suspended with 33 µL of anhydrous ACN. Seven microliters of TMTPro reagent in ACN was added to 9 µL of clean peptide in 0.1M TEAB. The final ACN concentration was 43.75% and the ratio of peptides to TMTPro reagent was 1:20. After 60 min of incubation the reaction was quenched with 2 µL of 5% hydroxylamine. Labelled peptides were pooled, dried, re-suspended in 200 µL of 0.1% formic acid (FA), split into two equal parts, and desalted using home-made STAGE tips. One of the two parts was fractionated on a 1 mm x 150 mm ACQUITY column, packed with 130 Å, 1.7 µm C18 particles (Waters cat. no SKU: 186006935), using an Ultimate 3000 UHPLC (Thermo Fisher Scientific). Peptides were separated at a flow of 30 µL/min with a 88 min segmented gradient from 1% to 50% buffer B for 85 min and from 50% to 95% buffer B for 3 min; buffer A was 5% ACN, 10mM ammonium bicarbonate (ABC), buffer B was 80% ACN, 10mM ABC. Fractions were collected every three minutes, pooled in two passes (1 + 17, 2 + 18 … etc.), and dried in a vacuum centrifuge (Eppendorf).

#### LC-MS/MS analysis

For label-free quantification, peptides were separated on a 75 cm, 75 μm internal diameter Acclaim™ PepMap™ analytical column (Thermo Fisher Scientific, catalogue number 164939) using an EASY-nLC 1200 (Thermo Fisher Scientific). The column was maintained at 50°C. Buffer A and B were 0.1% formic acid in water and 0.1% formic acid in 80% acetonitrile. Peptides derived from whole cells were separated on a segmented gradient from 6% to 31% buffer B for 230 min and from 31% to 50% buffer B for 10 min at 250 nl / min. Peptides derived from enriched mitochondria were separated on a segmented gradient from 6% to 31% buffer B for 110 min and from 31% to 50% buffer B for 10 min at 250 nl / min. Eluting peptides were analyzed on a QExactive HF mass spectrometer (Thermo Fisher Scientific). Peptide precursor m/z measurements were carried out at 60000 resolution in the 300 to 1800 m/z range. The top ten most intense precursors with charge state from 2 to 7 only were selected for HCD fragmentation using 25% normalized collision energy. The m/z values of the peptide fragments were measured at 30000 resolution using a minimum AGC target of 1e4 and 55 ms maximum injection time for the whole cell samples or 15000 resolution, minimum AGS target of 1e4, and 120 ms maximum injection time for the enriched mitochondria samples. Upon fragmentation, precursors were put on a dynamic exclusion list for 45 sec. TMT labeled peptides were and separated on a 50 cm, 75 µm Acclaim PepMap column (Thermo Fisher Scientific, Product No. 164942) and analysed on a Orbitrap Lumos Tribrid mass spectrometer (Thermo Fisher Scientific) equipped with a FAIMS device (Thermo Fisher Scientific). The FAIMS device was operated in two compensation voltages, −50 V and −70 V. Synchronous precursor selection based MS3 was used for the acquisition of the TMTPro reporter ion signals. Peptide separations were performed on an EASY-nLC1200 using a 90 min linear gradient from 6% to 31% buffer; buffer A was 0.1% FA, buffer B was 0.1% FA, 80% ACN. The analytical column was operated at 50°C. Raw files were split based on the FAIMS compensation voltage using FreeStyle (Thermo Fisher Scientific).

#### Protein identification and quantification

Label-free quantification raw data were analyzed with MaxQuant version 1.6.0.13 (50) using the integrated Andromeda search engine (51). Peptide fragmentation spectra were searched against the canonical sequences of the human reference proteome, proteome ID UP000000589, and toxoplasma reference proteome, proteome ID UP000005641, downloaded from UniProt. Methionine oxidation and protein N-terminal acetylation were set as variable modifications; cysteine carbamidomethylation was set as fixed modification. The digestion parameters were set to “specific” and “Trypsin/P,” The minimum number of peptides and razor peptides for protein identification was 1; the minimum number of unique peptides was 0. Protein identification was performed at a peptide spectrum matches and protein false discovery rate of 0.01. The “second peptide” option was on. Successful identifications were transferred between the different raw files using the “Match between runs” option. Exploratory data analysis and visualization was done using tidyverse in R (52, 53). TMT data was analyzed using MaxQuant, version 1.6.17.0. The isotope purity correction factors, provided by the manufacturer, were included in the analysis. Differential expression analysis was performed using limma, in R (54). Volcano plots of WC and mitoIP proteins generated using Flaski (55).

#### Stable Labelling with L-glutamine-^15^N_2_

2 million ES-2 cells were seeded in a 10cm dishes in DMEM containing 10% FBS and cultured for 16 h. Next day cells were washed once with 1x PBS, and the medium was replaced with glutamine-free DMEM containing 10% dFBS and 2 mM L-glutamine-^15^N_2_ (Cambridge isotope laboratories #NLM-1328-0.25) for 24h. The following day, cells were either mock-infected or infected with Toxoplasma (RHΔku80:mCherry+) at a multiplicity of infection (MOI) of 4 in cDMEM media in the absence of labelled L-glutamine-^15^N_2_ for 24 hpi. Parasites were isolated from infected monolayers and processed for metabolomics analysis.

#### Stable Labelling with 2,3,3-^2^H-serine

WT and ATF4 KO ES-2 cells (250,000) were seeded in DMEM containing 10% FBS and cultured for 16 h. The following day cells were washed once with PBS, and either mock-infected or infected with Toxoplasma strains RHΔku80:mCherry and RHΔcpsII:mCherry at a multiplicity of infection (MOI) of 4 and 6, respectively due to the lack of proliferation of RHΔcpsII:mCherry parasites in minimal essential media (MEM) containing 10% dialyzed FBS, 2 mM L-glutamine and 0.2 mM [2,3,3-^2^H] serine isotopologue (Cambridge Isotope Laboratories #DLM-1073-1). At 24 hpi, cells were harvested for metabolomics analysis.

#### Toxoplasma isolation for metabolite extraction

4 million cells ES-2 cells were seeded in 10 cm dishes overnight and infected with Toxoplasma at an MOI of 4 the subsequent morning. At 24 hpi, infected cells were collected by scraping and washed with ice-cold DPBS twice. All subsequent steps were performed at 4°C. Infected cells were resuspended in DPBS with phosphatase and proteases inhibitors and homogenized by passing 10 times through a 271/2 G needle. The homogenates were spun down at 1500 rpm for 5 min to remove cell debris and filtered with a 5 μM filter. The final pellets containing parasite-enriched fractions were resuspended in cold metabolite extraction buffer. The purity of the Toxoplasma-enriched fractions was assessed by SDS-PAGE and immunoblot.

#### Extraction of polar metabolites

Cells were washed twice with 75 mM of ammonium carbonate pH = 7.4 and the plates were frozen in liquid nitrogen and stored at −80°C until metabolite extraction. For metabolite extraction, 1ml of −20°C cold extraction buffer (HPLC-grade ultrapure 40% MeOH, 40% acetonitrile, 20% water, containing 250 nM ^13^C^15^N Amino acids (Cambridge isotopes MSK_A2-1.2), 100 ng/mL of ^13^C_10_ ATP (Sigma 710695), 100 ng/mL ^15^N_5_ ADP (Sigma 741167), 100 ng/mL ^13^C_10_^15^N_5_ AMP (Sigma 650676) and 20 ng/mL citric acid ^2^H_4_ (Sigma 741167) as isotope-labeled internal standards.) was added to each well of the frozen plate. The cells were then scraped, transferred to a new tube and dissolved by sonication followed by an incubation for 30 min at 1500 rpm at 4°C. The metabolite-containing supernatant was cleared by centrifugation at 20,000 × g for 10 min and subsequently transferred to a speedvac concentrator to fully evaporate the extraction buffer and stored at −80°C until further analysis. The remaining protein pellet from the extraction was used to determine protein concentration of the sample.

#### Anion-Exchange Chromatography Mass Spectrometry (AEX-MS) for the analysis of anionic metabolites

Extracted metabolites were re-suspended in 150 µl of UPLC/MS grade water (Biosolve), of which 100 µl were transferred to polypropylene autosampler vials (Chromatography Accessories Trott, Germany) before AEX-MS analysis. The samples were analysed using a Dionex ionchromatography system (Integrion Thermo Fisher Scientific) as described previously (56). In brief, 5 µL of the resuspended polar metabolite extract were injected in push-partial mode, using an overfill factor of 1, onto a Dionex IonPac AS11-HC column (2 mm × 250 mm, 4 μm particle size, Thermo Fisher Scientific) equipped with a Dionex IonPac AG11-HC guard column (2 mm × 50 mm, 4 μm, Thermo Fisher Scientific). The column temperature was held at 30°C, while the auto sampler temperature was set to 6°C. A potassium hydroxide gradient was generated using a potassium hydroxide cartridge (Eluent Generator, Thermo Scientific), which was supplied with deionized water (Milli-Q IQ 7000, Millipore). The metabolite separation was carried at a flow rate of 380 µL/min, applying the following gradient conditions: 0-3 min, 10 mM KOH; 3-12 min, 10−50 mM KOH; 12-19 min, 50-100 mM KOH; 19-22 min, 100 mM KOH, 22-23 min, 100-10 mM KOH. The column was re-equilibrated at 10 mM for 3 min. For the analysis of metabolic pool sizes the eluting compounds were detected in negative ion mode using full scan measurements in the mass range m/z 77 – 770 on a Q-Exactive HF high resolution MS (Thermo Fisher Scientific). The heated electrospray ionization (ESI) source settings of the mass spectrometer were: Spray voltage 3.2 kV, capillary temperature was set to 300°C, sheath gas flow 50 AU (arbitrary units), aux gas flow 20 AU at a temperature of 330°C and a sweep gas flow of 2 AU. The S-lens was set to a value of 60. The LC-MS data analysis was performed using the TraceFinder software (Version 5.1, Thermo Fisher Scientific). The identity of each compound was validated by authentic reference compounds, which were measured at the beginning and the end of the sequence. For data analysis the area of the deprotonated [M-H^+^]^-1^ or doubly deprotonated [M-^2^H]^-2^ isotopologue mass peaks of every required compound were extracted and integrated using a mass accuracy <3 ppm and a retention time (RT) tolerance of <0.05 min as compared to the independently measured reference compounds. If no independent ^12^C experiments were carried out, where the pool size is determined from the obtained peak area of the ^12^C monoisotopologue, the pool size determination was carried out by summing up the peak areas of all detectable isotopologues per compound. These areas were then normalized, as performed for un-traced ^12^C experiments, to the internal standards, which were added to the extraction buffer, followed by a normalization to the protein content or the cell number of the analyzed samples. The relative isotope distribution of each isotopologue was calculated from the proportion of the peak area of each isotopologue towards the sum of all detectable isotopologues. Natural abundance of the measured isotopologuge was corrected using the AccuCor package (57).

#### Sample preparation for folate detection

Cell extracts were prepared by homogenization of cell pellets by sonication in buffer containing 20 mM ammonia acetate, 0.1% ascorbic acid, 0.1% citric acid and 100 mM dithiothreitol at pH 7. Protein was removed by precipitation by addition of two volumes of acetonitrile and centrifugation (12,000 g at 4°C). Supernatants were transferred, lyophilized, stored at −80°C and re-suspended in dH_2_O before analysis. Folate analysis was carried out by ultraperformance liquid chromatography tandem mass spectrometry (UPLC-MS/MS) as described previously (58, 59). Lyophilized samples were resuspended in 25 uL water (milli-Q) and centrifuged for 5 min at 12,000 g at 4°C. Metabolites were resolved by reversed-phase UPLC (Acquity UPLC BEH C18 column, Waters Corporation, UK). Solvents for UPLC were as follows: Buffer A, 5% methanol, 95% Milli-Q water and 5 mM dimethylhexylamine at pH 8.0; Buffer B, 100% methanol, 5 mM dimethylhexylamine. The column was equilibrated with 95% Buffer A: 5% Buffer B. The sample injection volume was 20 uL. The UPLC protocol consisted of 95% Buffer A: 5% Buffer B for 1 min, followed by a gradient of 5–60% Buffer B over 9 min and then 100% Buffer B for 6 min before re-equilibration for 4 min. The metabolites were eluted at a flow rate of 500 μL/min and the wash step with 100% Buffer B was at flow rate of 600 μL/min. The UPLC was coupled to a XEVO-TQS mass spectrometer (Waters Corporation) operating in negative-ion mode using the following settings: capillary 2.5 kV, source temperature 150°C, desolvation temperature 600°C, cone gas flow rate 150 L/h, and desolvation gas flow rate 1200 L/h. Folates were measured by multiple reaction monitoring with optimized cone voltage and collision energy for precursor and product ions (58, 60). Peak areas were analysed by TargetLynx software (Waters Corporation, UK).

#### Infection of mice with Toxoplasma

8 week old female C57BL/6 J mice were obtained from Charles River. All animal work was approved by local authorities (Landesamt für Natur, Umwelt und Verbraucherschutz Nordrhein-Westfalen, Germany) and animal procedures were carried out in accordance with European, national and institutional guidelines and according to good practice of animal handling. Mice were maintained at the Max Planck Institute for Biology of Ageing with 12 h light cycle and regular chow diet. When mice were 10 weeks old, following serial dilution in PBS, RHΔku80:mCherry tachyzoites were injected intraperitoneally. Tunicamycin (Sigma, #11089-65-9) was dissolved in 150 mM sucrose in PBS and injected intraperitoneally at 1 mg kg^−1^as previously described (61, 62). ISRIB (Sigma, #1597403-47-8) was dissolved in DMSO at 6.25 mg ml^−1^ and subsequently diluted in PBS at 0.25 mg ml^−1^ and delivered intraperitoneally to mice at a dose of 2.5 mg kg^−1^ per day every other day in the morning as previously described in (63). For vehicle DMSO was diluted in PBS.

#### PEC isolation

Euthanized C57BL/6 J mice (n = 5 per treatment) were injected with 6 ml cold 1× PBS using a 26-gauge needle. Mice were palpated for 1–2 min, following which fluid contents were aspirated out of the peritoneum. Content was spun down at 1,000 rpm for 5 min, washed, and resuspended in DMEM to a concentration of 2×10^6^ per ml. Samples were then pellet and snap frozen in liquid nitrogen until further downstream analysis was performed accordingly.

#### Statistical analysis

All statistical analyses were performed using one-way ANOVA, two-way ANOVA, an unpaired t-test or multiple unpaired t-tests in GraphPad Prism 9 software and are indicated accordingly.

---

**Fig. S1.**
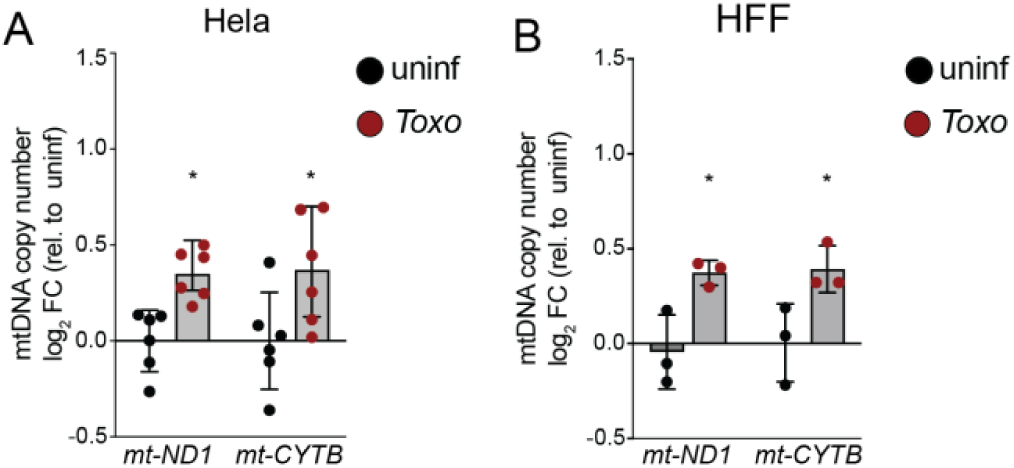
*Toxoplasma* infection drives an increase in mtDNA copy number in both cancer and primary cell lines. mtDNA levels assessed by qPCR for *mt-ND1* and *mt-CYTB* (normalized to *RUNX2*) in (**A**) HeLa cells and (**B**) primary human foreskin fibroblasts (HFFs) that were uninfected or infected with *Toxoplasma* at a multiplicity of infection (MOI) of 4 and harvested at 24 hours post infection (hpi). Data are mean ± SD of (**A**) n=6 independent cultures and (**B**) n=3 independent cultures; *p<0.05; for uninfected versus infected by means of t-tests analysis.

**Fig. S2.**
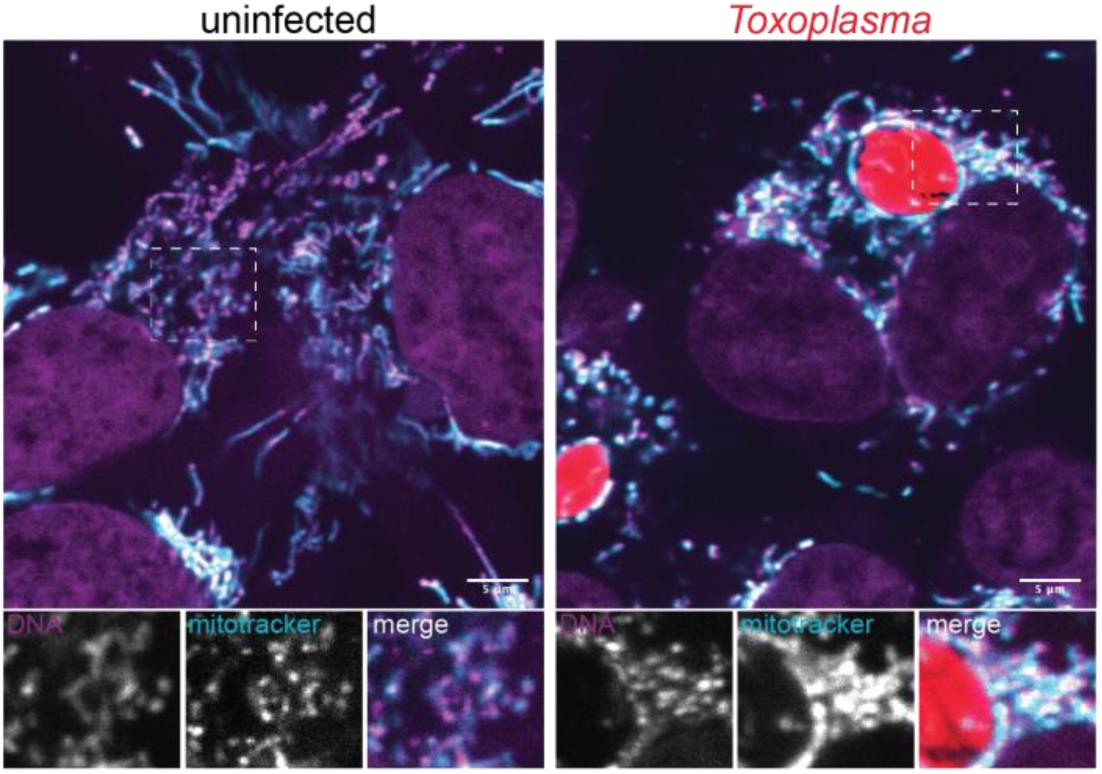
*Toxoplasma* infection does not induce mtDNA release into the cytosol. Representative live-cell images of uninfected and *Toxoplasma-*mCherry-infected ES-2 cells labeled with MitoTracker Deep Red (mitotracker) and the DNA dye picogreen (DNA) and imaged at 24 hpi. Scale bar, 5 µm.

**Fig. S3.**
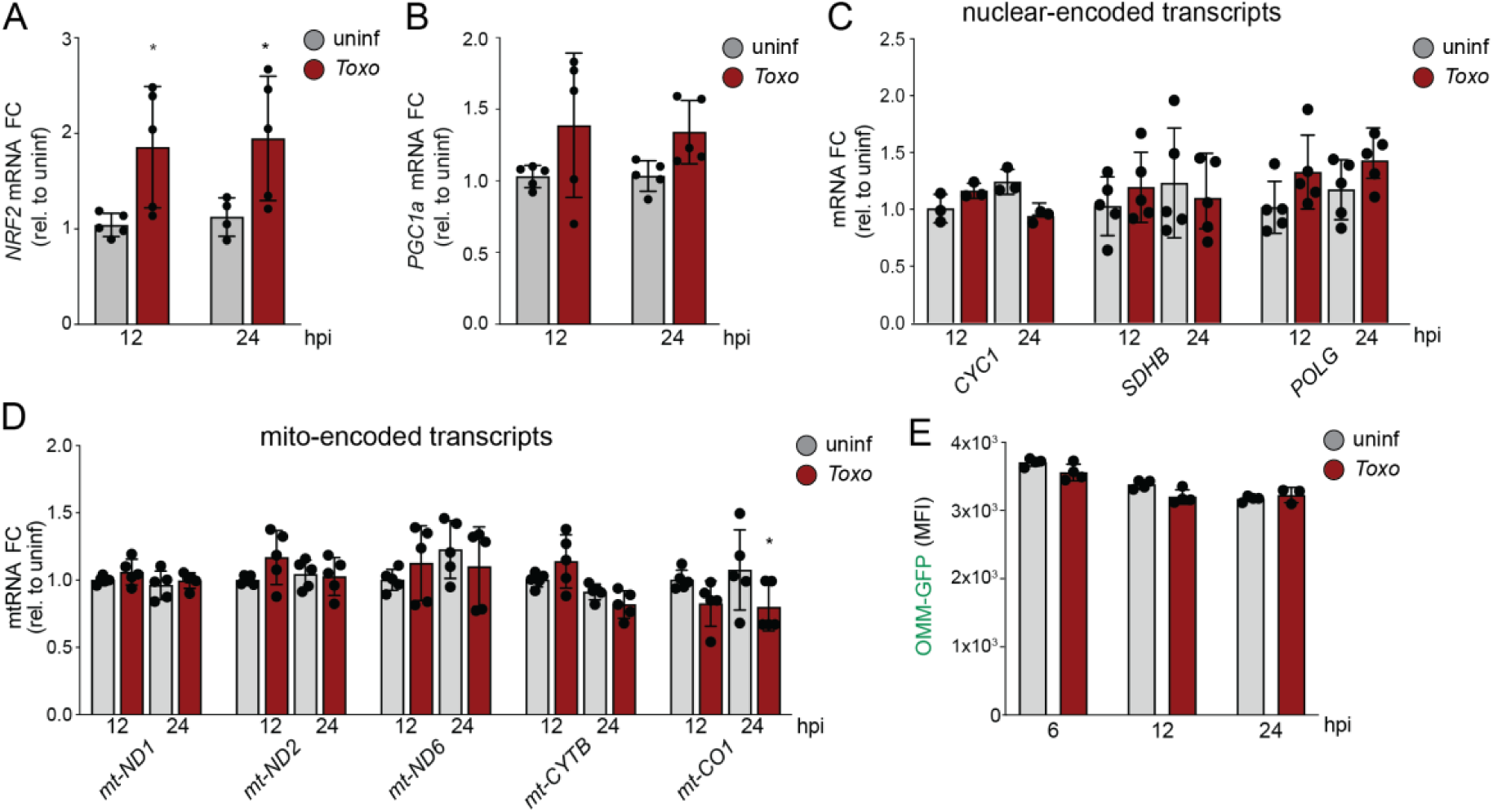
*Toxoplasma* infection does not induce mitochondrial biogenesis. Uninfected and *Toxoplasma*-infected cells (MOI=4) were analyzed by qPCR at the indicated timepoints for (**A**) *NRF2*; (**B**) PGC1α; (**C**) the nuclear-encoded mitochondrial transcripts *CYC1*, *SDHB*, *POLG;* and (**D**) the mitochondrial-encoded transcripts *mt-ND1, mt-ND2, mt-ND6, mt-CYTB and mt-CO1.* Transcripts were normalized to *ACTB* and are relative to uninfected samples. Data are mean ± SD of n=5 independent cultures, *p<0.05; for uninfected versus infected by means of two-way ANOVA analysis. (**E**) GFP fluorescence intensity (MFI) determined by flow cytometry analysis of cells stably expressing GFP targeted to the outer mitochondrial membrane (OMM) in uninfected cells or cells infected with RH-mCherry expressing *Toxoplasma* at 6, 12 and 24 hpi. Data are mean ± SEM of n=3 independent cultures. ES-2 cells used for all experiments.

**Fig. S4.**
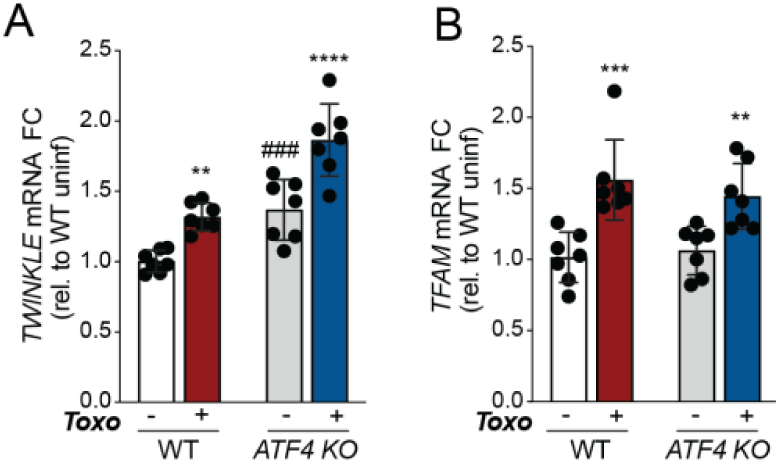
*TWNK* and *TFAM* are induced independently of ATF4 during *Toxoplasma* infection. Uninfected or *Toxo-*infected WT and *ATF4* KO ES-2 cells were analyzed by qPCR for (**A**) *TWINKLE* and (**B**) *TFAM* at 24 hpi. (**A** and **B**) Transcript levels were normalized to *ACTB* and are relative to WT uninfected. Data are mean ± SD of n=7 independent cultures. **p < 0.01; ***p<0.001; ****p<0.001 for uninfected versus *Toxo*-infected, ###*p* < 0.001 for WT versus *ATF4 KO* by means of two-way ANOVA analysis.

**Fig. S5.**
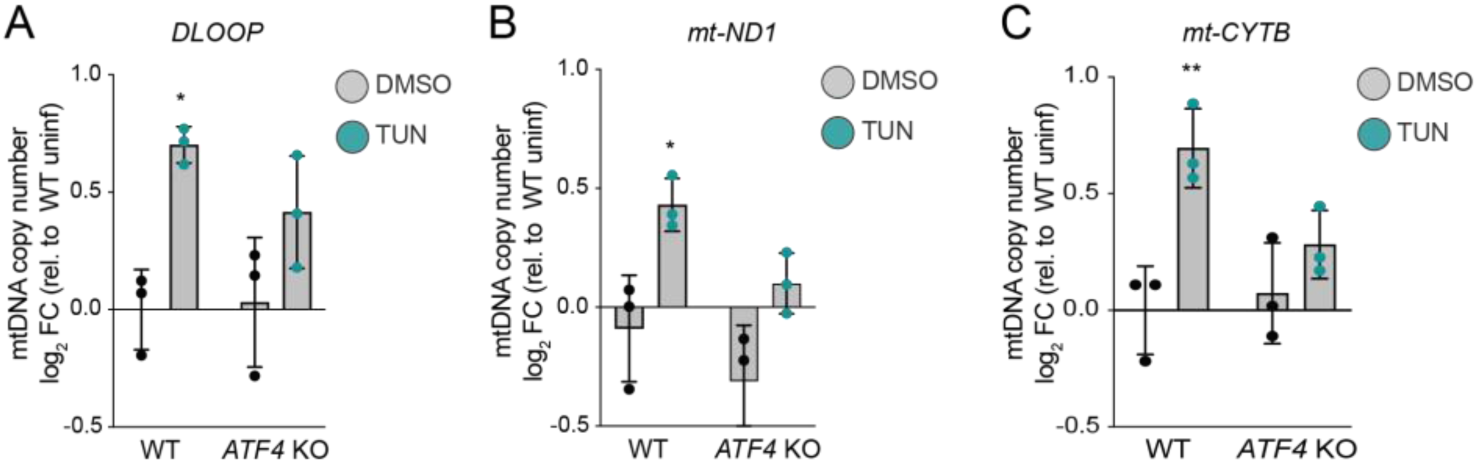
ISR activation increases mtDNA levels independently of infection. mtDNA levels were assessed by qPCR of (**A**) *DLOOP,* (**B**) *mt-ND1* and (**C**) *mt-CYTB* (normalized to *RUNX2*) in WT and *ATF4* KO ES-2 cells that were untreated or treated with tunicamycin (3 μg/ml) for 16h. Data are mean ± SD of n=3 independent cultures; *p<0.05; **p<0.001; for untreated versus tunicamycin-treated by means of two-way ANOVA analysis.

**Fig. S6.**
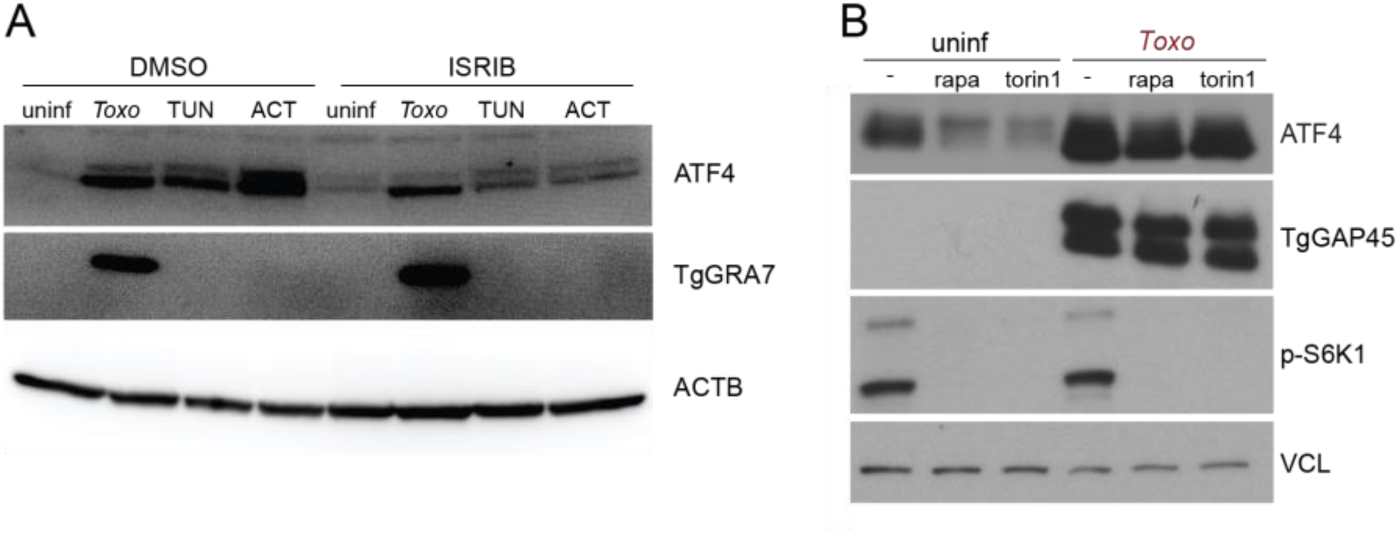
ATF4 activation during infection is mTOR independent. (**A**) Immunoblot (IB) analysis of lysates from ES-2 cells that were treated as indicated: uninfected (uninf); *Toxoplasma (Toxo)* at MOI=4; tunicamycin (TUN; 3 μg/ml); or actinonin (ACT; 50 μM) with or without with ISRIB (200 nM) for 24h: ATF4, ∼50 kDa; α-Actin (ACTB), ∼35 kDa and *Toxoplasma* GRA7 (TgGRA7), ∼27 kDa. (**B**) Immunoblot analyses of lysates from uninfected and *Toxo-*infected (MOI: 4) ES-2 cells at 24 hpi and co-treated with DMSO, rapamycin (rapa; 20 nM) or torin1 (250 nM) and probed with the following antibodies: ATF4, ∼50 kDa; Phospho-p70 S6 Kinase Thr389 (p-S6K1), ∼70 and 80 kDa; Vinculin (VCL), ∼124 kDa; and *Toxoplasma* GAP45 (TgGAP45,) ∼45 kDa.

**Fig. S7.**
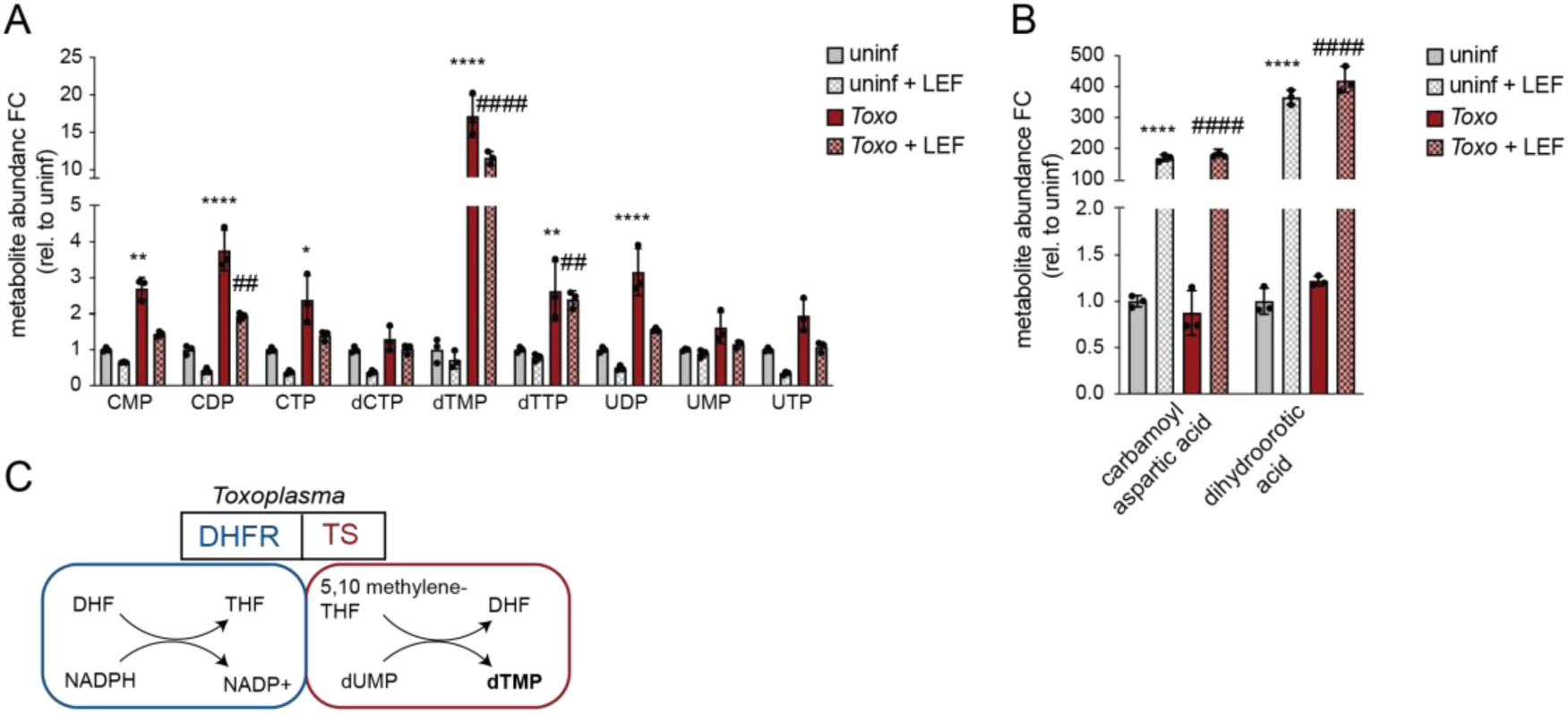
dTMP levels are increased in *Toxoplasma-*infected cells during host pyrimidine synthesis inhibition. (**A**) Abundance of total pyrimidine intermediates metabolites in ES-2 uninfected and *Toxoplasma*-infected cells at MOI=4 and treated with DMSO or leflunomide (LEF) (50 μM). Data are mean ±SEM of n=3 independent cultures, and are normalized to cell number, **p<0.01; ****p<0.0001 for uninf versus infected and ##p<0.01; ####p < 0.0001 for DMSO versus LEF treatment by means of two-way ANOVA analysis. (**B**) Total abundance of carbamoyl aspartic acid and dihydroorotic acid in samples treated as in (A). Data are mean ±SEM of n=3 independent cultures, and are normalized by cell number, ****p<0.0001 for uninfected versus infected and ####p < 0.0001 for DMSO versus LEF treatment by means of two-way ANOVA analysis. (**C**) Schematic of bifunctional dihydrofolate reductase thymidylate synthase (DHFR-TS) of *Toxoplasma gondii*.

**Fig. S8.**
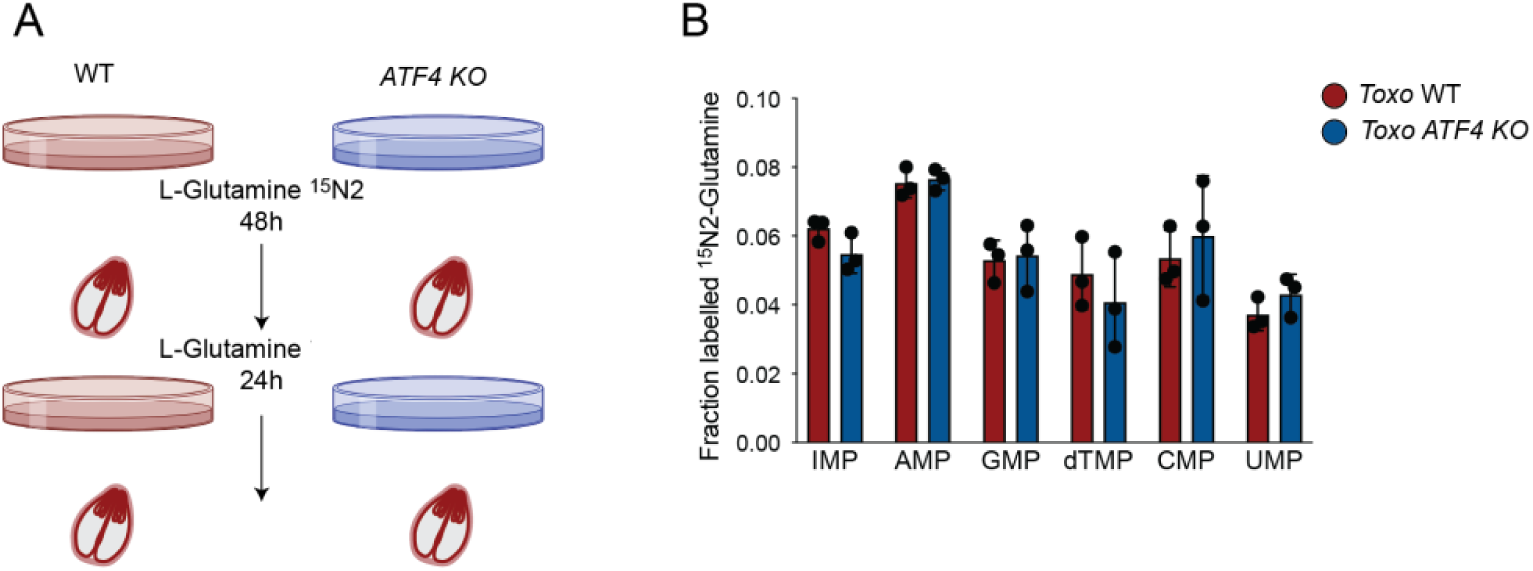
Loss of ATF4 does not promote nucleotide acquisition by *Toxoplasma*. (**A**) Workflow for assessing *Toxoplasma* nucleotide acquisition from host cells. WT and *ATF4* KO ES-2 cells were culture with 2 mM L-glutamine^15^N_2_ for 48 h. Cells were then infected with *Toxoplasma* in the absence of the isotopologue L-glutamine^15^N_2_. After 24h the *Toxoplasma*-enriched fraction was pelleted for metabolite extraction. (**B**) Fraction enrichment of glutamine-derived ^15^N nucleotides of *Toxoplasma*-purified extracts from WT and *ATF4* KO cells as described in (A). Data are ± SD of n=3 independent cultures.

**Fig. S9.**
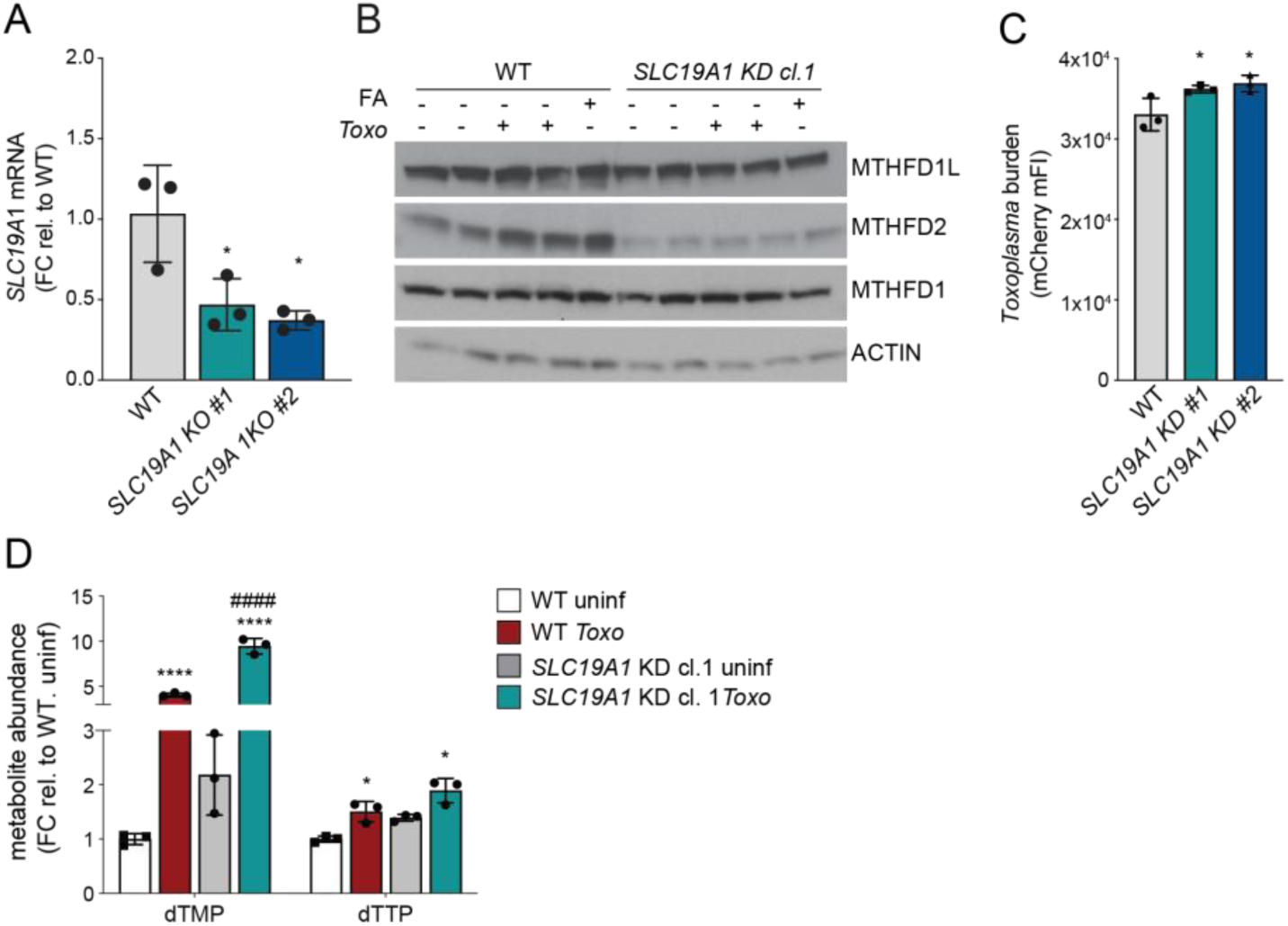
Increased parasite proliferation and dTMP levels in a model of mitochondrial 1C-dysfunction. (**A**) WT and SLC19A KO cells were analyzed by qPCR for *SLC19A1*. Transcripts were normalized to *ACTB*. Data are mean ± SD of n=3 independent cultures. (**B**) WT and *SLC19A1* knockdown (KD) ES-2 cells were cultured for 4 days in RPMI medium lacking folic acid and uninfected, infected with *Toxoplasma* (MOI: 4) cells or supplemented with folic acid (2 μM). After 24h, samples were analyzed by immunoblotting for the indicated antibodies MTHFD1L, ∼106 kDa; MTHFD2, ∼32 kDa; MTHFD1, ∼40 kDa and α-Actin (ACTB), ∼35 kDa. (**C**) WT and S*LC19A1* KD ES-2 cells were infected with *Toxoplasma* and analyzed 24 hpi by means of flow cytometry for *Toxoplasma* burden [mCherry median FI]. Data are mean ± SEM of three biological experiments, *p < 0.05 for WT versus *SLC19A1* KO, by means of one-way ANOVA analysis. (**D**) Total abundance of dTMP and dTTP of WT and S*LC19A1* KO ES-2 uninfected and *Toxo*-infected cells. Data are mean ± of SED n=3 independent cultures, and normalized by cell number, *p<0.05; ****p<0.0001 for uninfected versus infected cells and ####p < 0.0001 for WT versus *SLC19A1* KO cells by means of two-way ANOVA analysis.

**Fig. S10.**
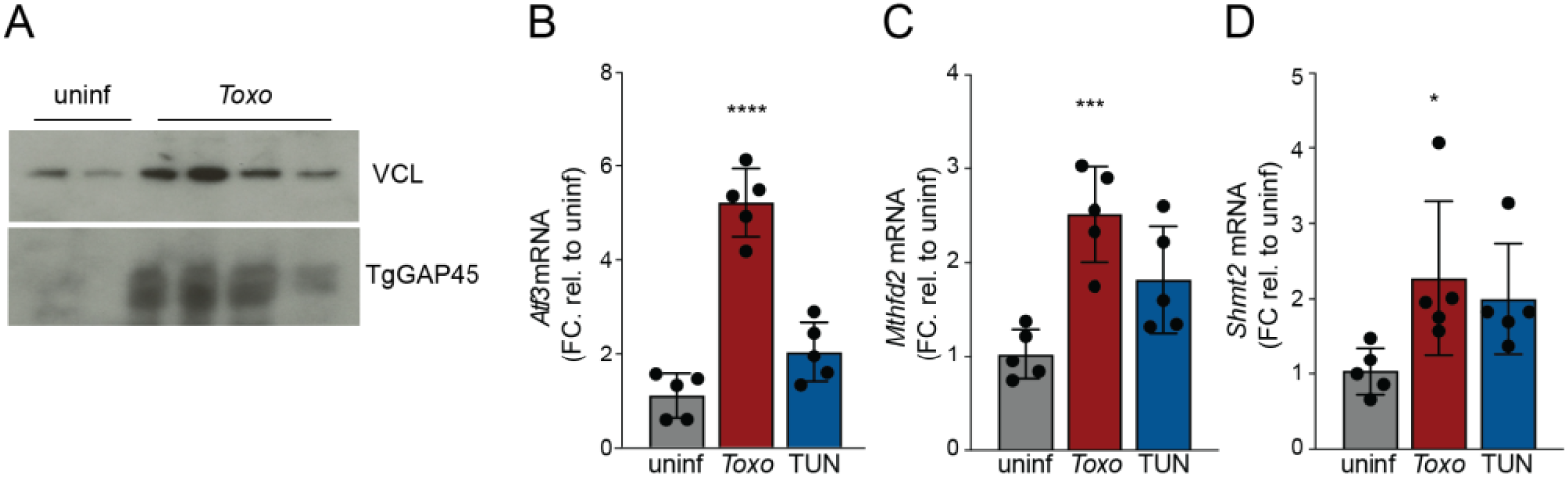
*Toxoplasma* infection and tunicamycin treatment induce ATF4 targets in vivo. (**A**) Immunoblot (IB) analysis of lysates of peritoneal exudate cells (PECs) isolated at 5 dpi from mice that there uninfected or infected with 50 tachyzoites: *Toxoplasma* GAP45 (TgGAP45), ∼45 kDa; Vinculin (VCL), ∼124 kDa; (**B**-**D**) Mice were injected intraperitoneally with 1x PBS or 50 tachyzoites in 1x PBS or with tunicamycin (1 mg kg^−1^ body weight). At 5 days post injection (dpi) peritoneal exudate cells (PECs) were isolated and analyzed for the indicated ATF4 targets genes by qPCR. Transcripts are normalized to *Hprt* and relative to untreated uninfected PECs mice (n=5). * p<0.05; *** p<0.001; and ****p<0.0001 by means of one-way ANOVA analysis.

