## supplementary file for "Metabolic immunity to infection is driven by mitochondrial one-carbon metabolism"

**Affiliations:**

**This PDF file includes:**

Materials and Methods

Figs. S1 to S10

References 42-63

**Other Supplementary Materials for this manuscript include the following:**

Supplementary Tables S1-S3

**Materials and Methods**

*Cell culture and cell lines*
HeLa adenocarcinoma cells, ES-2 ovary clear cell carcinoma, Human foreskin fibroblasts (HFF), were obtained from ATCC (CCL-2, CRL-1978, and SCRC-1041, respectively); ES-2 cells stably expressing pMXs-eGFP-OMP25 (referred to as OMM-GFP; Addgene #83356) were generated through lentiviral transduction. Unless stated otherwise cell lines were cultured in Dulbecco's Modified Eagle's Medium (DMEM) containing 10% heat-inactivated fetal bovine serum (FBS) and maintained at 37°C and 5% CO_2_. AN3-12 mouse embryonic stem cells were obtained from Dr. Josef Penninger (*42*) and were cultured in DMEM high glucose supplemented with glutamine, 15% fetal bovine serum, penicillin/streptomycin, nonessential amino acids, sodium pyruvate, β-mercaptoethanol, and LIF as previously described (*43*). *ATF4* KO ES-2s were cultured in DMEM high glucose, 10% fetal bovine serum, β-mercaptoethanol and nonessential amino acids. Cells were tested every 2 weeks for *Mycoplasma* infection by means of polymerase chain reaction (PCR).

*Parasite culture and strains*

*Toxoplasma gondii* parasites of the Type I RHΔ*ku80:mCherry+* (previously described in (*44*)); Type I RHΔ*myr1* parasites and RHΔ*myr1::Myr1* were provided by Dr. Moritz Treeck, (Gulbenkian Institute) (*45*); Type I RHΔcpsII:mCherry parasites were provided by Dr. David Bzik (Dartmouth Geisel School of Medicine) (*46*). Parasites were maintained by serial passage in human foreskin fibroblast (HFF) monolayers in cDMEM. RHΔ*cpsII*: mCherry were maintained by serial passage in human foreskin fibroblast (HFF) monolayers in cDMEM supplemented with 0.2 mM uracil.

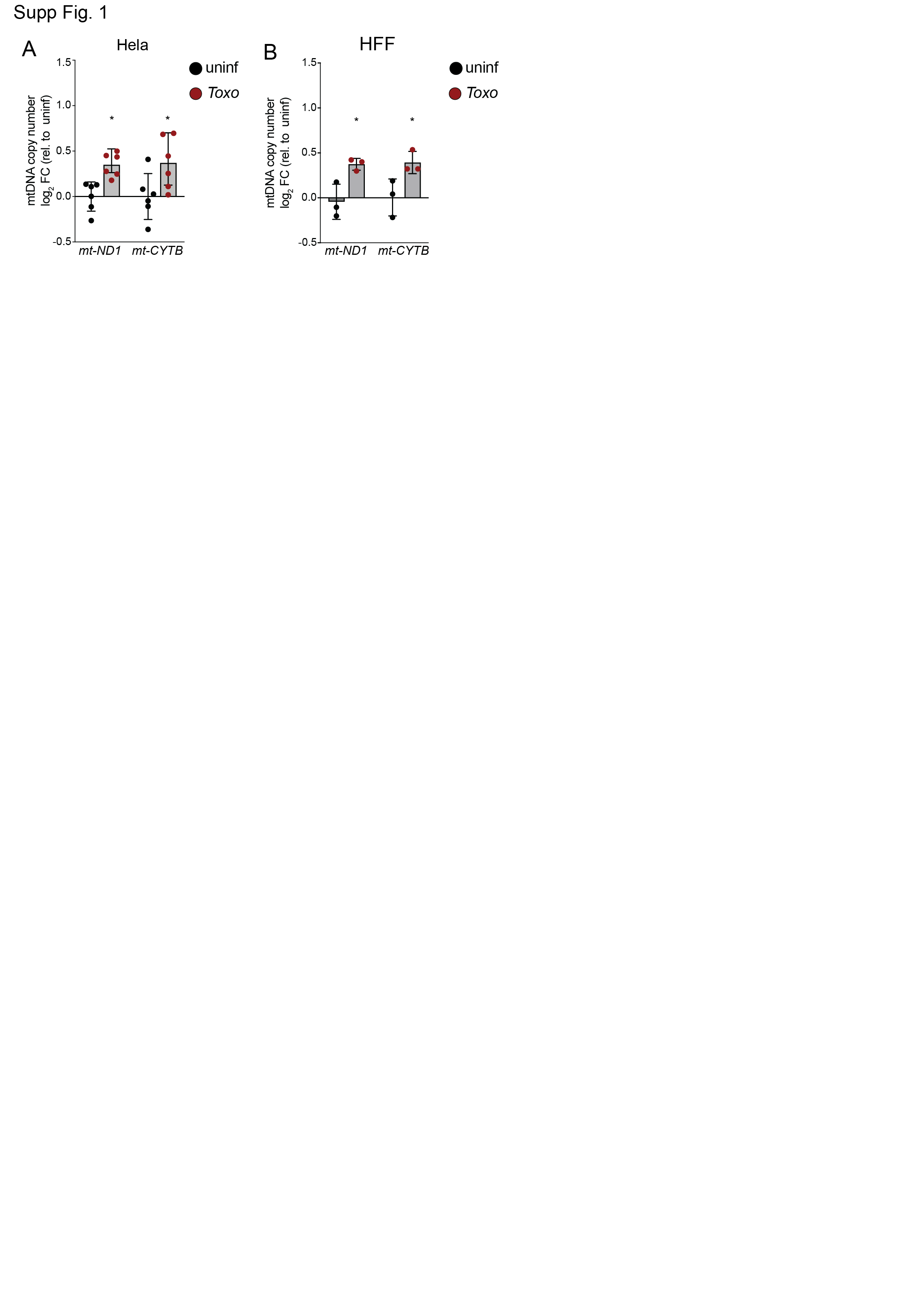

Fig. S1. *Toxoplasma* infection drives an increase in mtDNA copy number in both cancer and primary cell lines. mtDNA levels assessed by qPCR for *mt-ND1* and *mt-CYTB* (normalized to *RUNX2*) in (A) HeLa cells and (B) primary human foreskin fibroblasts (HFFs) that were uninfected or infected with *Toxoplasma* at a multiplicity of infection (MOI) of 4 and harvested at 24 hours post infection (hpi). Data are mean ± SD of (A) n=6 independent cultures and (B) n=3 independent cultures; *p<0.05; for uninfected versus infected by means of t-tests analysis.

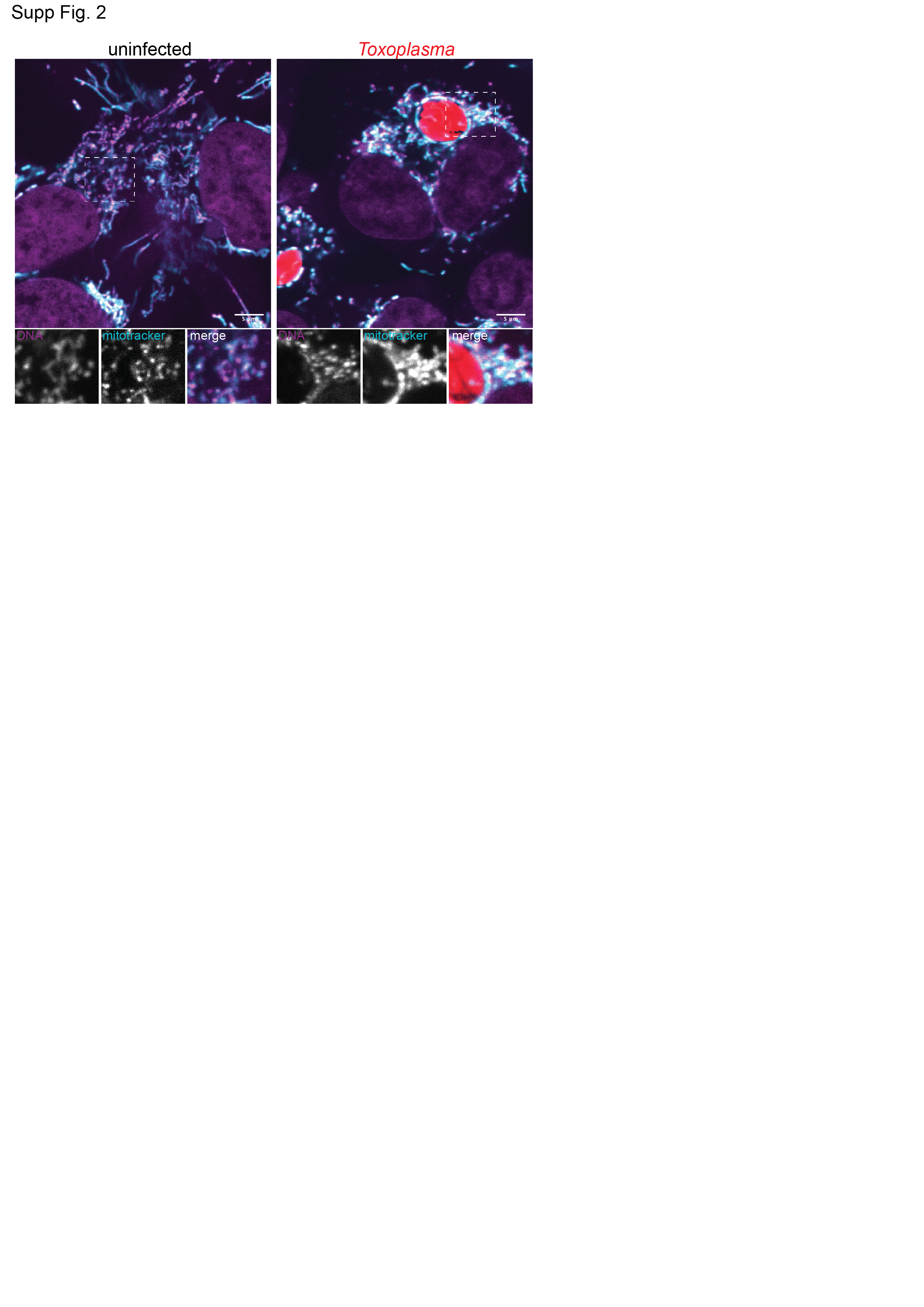
Fig. S2. *Toxoplasma* infection does not induce mtDNA release into the cytosol. Representative live-cell images of uninfected and *Toxoplasma-*mCherry-infected ES-2 cells labeled with MitoTracker Deep Red (mitotracker) and the DNA dye picogreen (DNA) and imaged at 24 hpi. Scale bar, 5 µm.

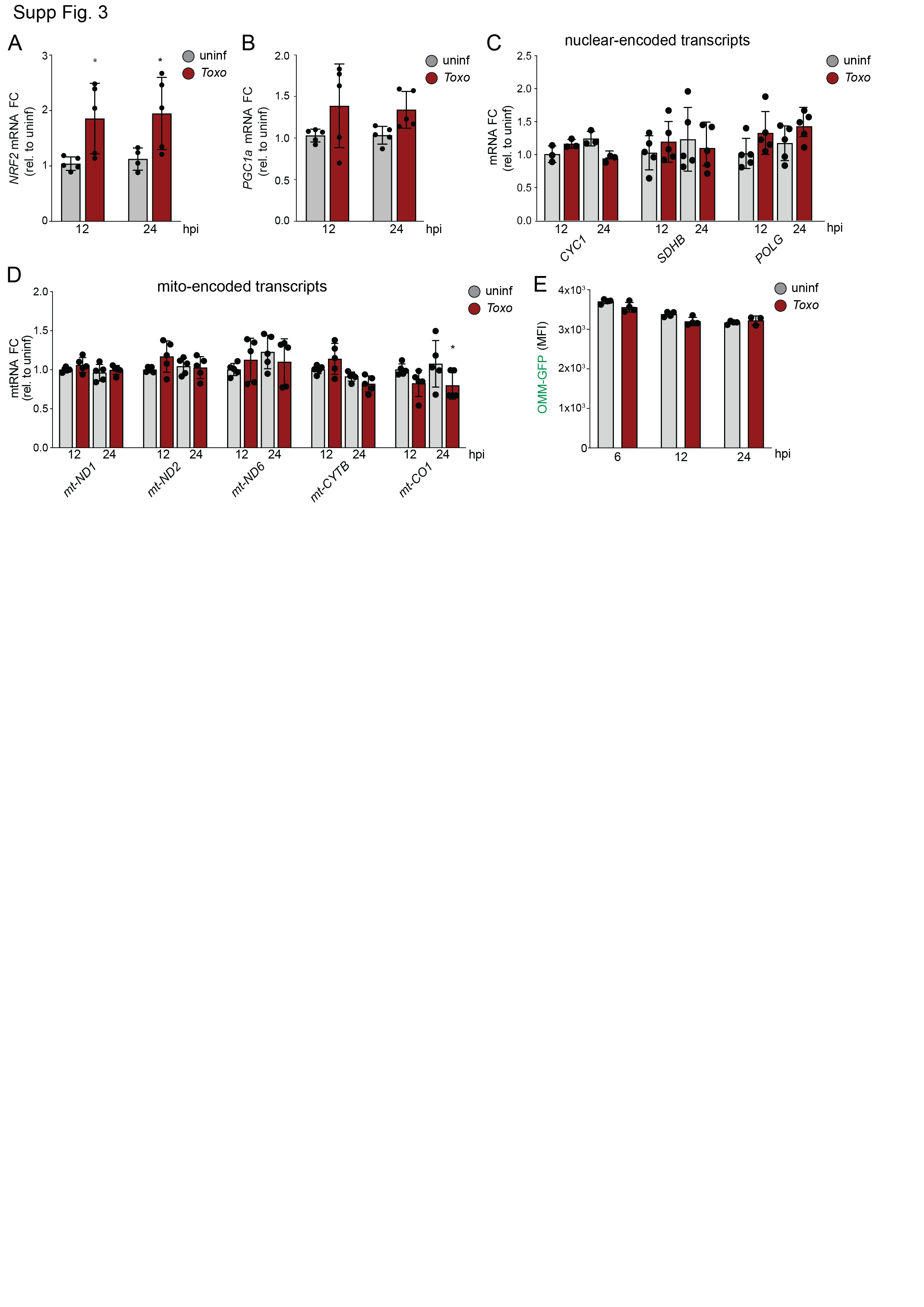

**Fig. S3. *Toxoplasma* infection does not induce mitochondrial biogenesis.** Uninfected and *Toxoplasma*-infected cells (MOI=4) were analyzed by qPCR at the indicated timepoints for (**A**) *NRF2*; (**B**) PGC1α; (**C**) the nuclear-encoded mitochondrial transcripts *CYC1*, *SDHB*, *POLG;* and (**D**) the mitochondrial-encoded transcripts *mt-ND1, mt-ND2, mt-ND6, mt-CYTB and mt-CO1.* Transcripts were normalized to *ACTB* and are relative to uninfected samples. Data are mean ± SD of n=5 independent cultures, *p<0.05; for uninfected versus infected by means of two-way ANOVA analysis. (**E**) GFP fluorescence intensity (MFI) determined by flow cytometry analysis of cells stably expressing GFP targeted to the outer mitochondrial membrane (OMM) in uninfected cells or cells infected with RH-mCherry expressing *Toxoplasma* at 6, 12 and 24 hpi. Data are mean ± SEM of n=3 independent cultures. ES-2 cells used for all experiments.

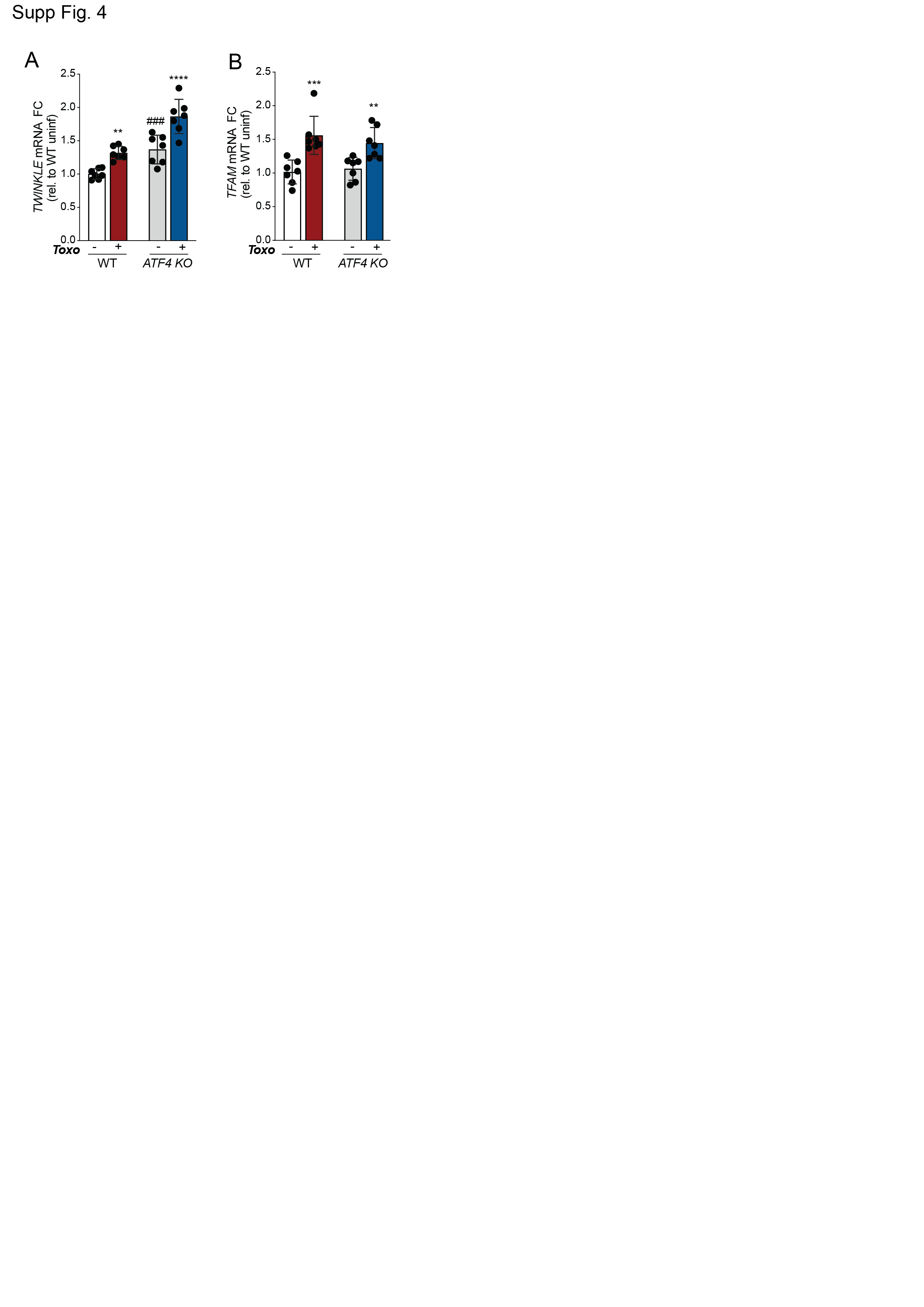
**Fig. S4. *TWNK* and *TFAM* are induced independently of ATF4 during *Toxoplasma* infection.** Uninfected or *Toxo-*infected WT and *ATF4* KO ES-2 cells were analyzed by qPCR for (**A**) *TWINKLE* and (**B**) *TFAM* at 24 hpi. (**A** and **B**) Transcript levels were normalized to *ACTB* and are relative to WT uninfected. Data are mean ± SD of n=7 independent cultures. **p < 0.01; ***p<0.001; ****p<0.001 for uninfected versus *Toxo*-infected, ###*p* < 0.001 for WT versus *ATF4 KO* by means of two-way ANOVA analysis.

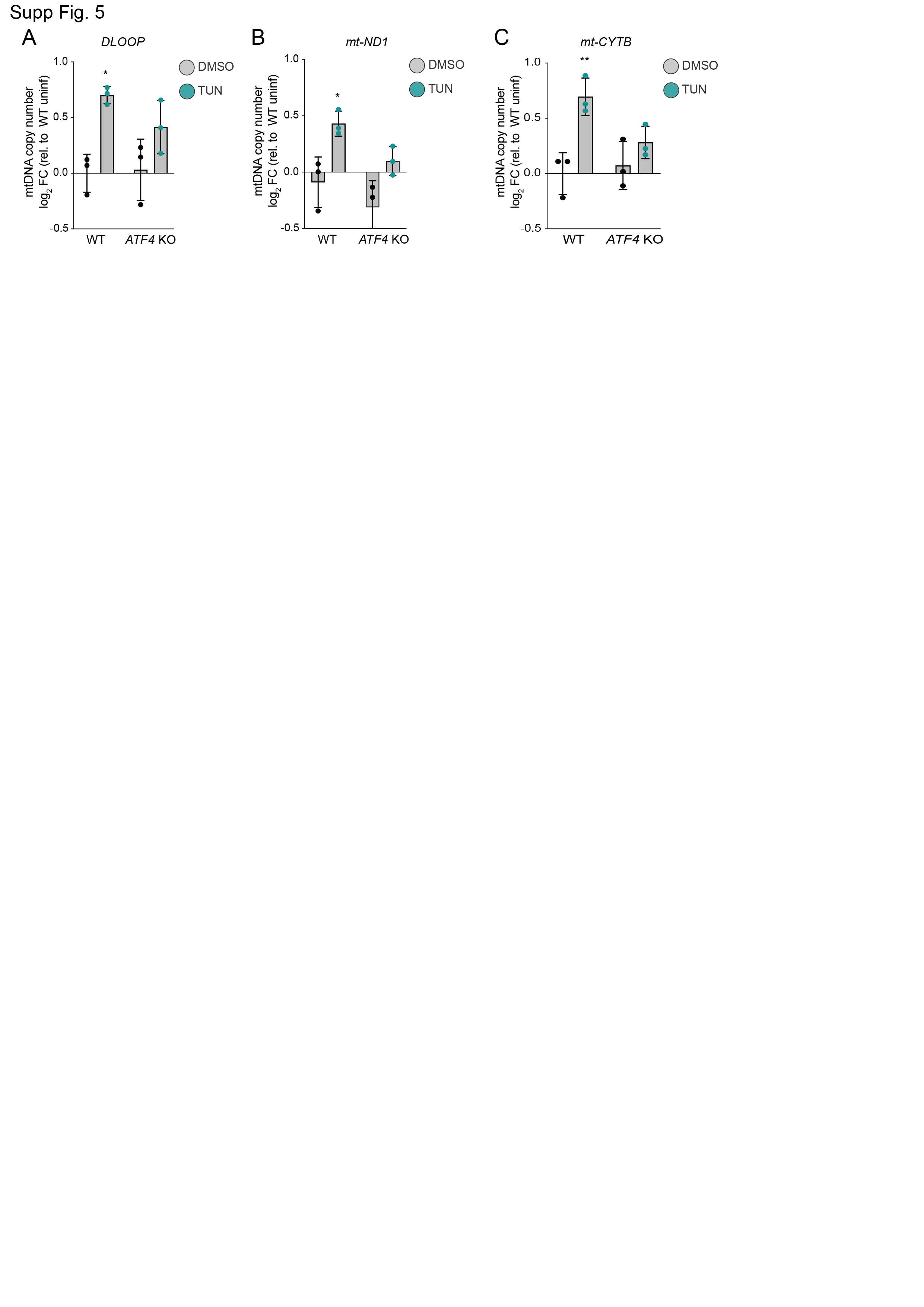

**Fig. S5. ISR activation increases mtDNA levels independently of infection.** mtDNA levels were assessed by qPCR of (**A**) *DLOOP,* (**B**) *mt-ND1* and (**C**) *mt-CYTB* (normalized to *RUNX2*) in WT and *ATF4* KO ES-2 cells that were untreated or treated with tunicamycin (3 μg/ml) for 16h. Data are mean ± SD of n=3 independent cultures; *p<0.05; **p<0.001; for untreated versus tunicamycin-treated by means of two-way ANOVA analysis.

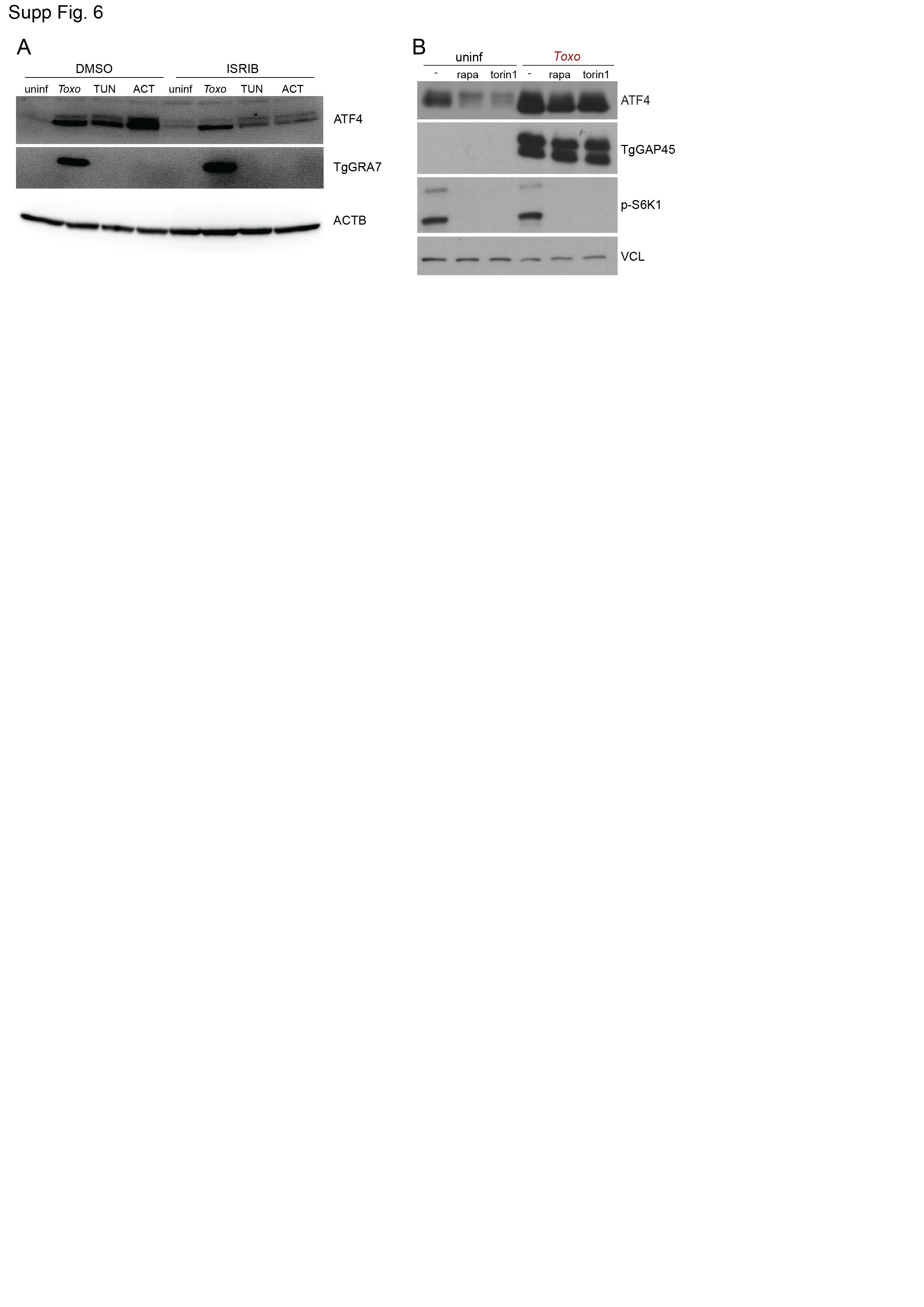
**Fig. S6. ATF4 activation during infection is mTOR independent.** (**A**) Immunoblot (IB) analysis of lysates from ES-2 cells that were treated as indicated: uninfected (uninf); *Toxoplasma (Toxo)* at MOI=4; tunicamycin (TUN; 3 μg/ml); or actinonin (ACT; 50 μM) with or without with ISRIB (200 nM) for 24h: ATF4, ~50 kDa; α-Actin (ACTB), ~35 kDa and *Toxoplasma* GRA7 (TgGRA7), ~27 kDa. (**B**) Immunoblot analyses of lysates from uninfected and *Toxo-*infected (MOI: 4) ES-2 cells at 24 hpi and co-treated with DMSO, rapamycin (rapa; 20 nM) or torin1 (250 nM) and probed with the following antibodies: ATF4, ~50 kDa; Phospho-p70 S6 Kinase Thr389 (p-S6K1), ~70 and 80 kDa; Vinculin (VCL), ~124 kDa; and *Toxoplasma* GAP45 (TgGAP45,) ~45 kDa.

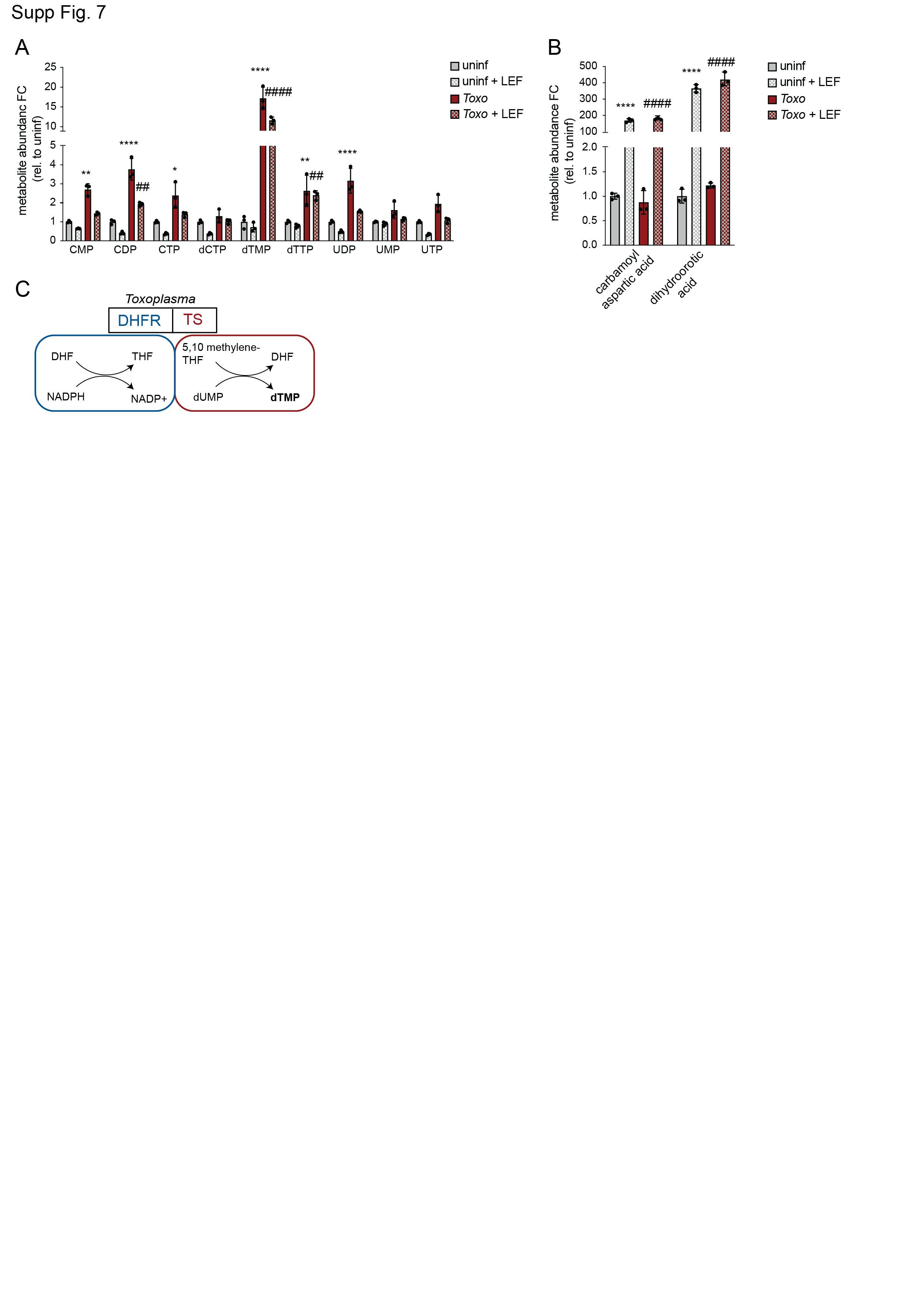
**Fig. S7. dTMP levels are increased in *Toxoplasma-*infected cells during host pyrimidine synthesis inhibition.** (**A**) Abundance of total pyrimidine intermediates metabolites in ES-2 uninfected and *Toxoplasma*-infected cells at MOI=4 and treated with DMSO or leflunomide (LEF) (50 μM). Data are mean ±SEM of n=3 independent cultures, and are normalized to cell number, **p<0.01; ****p<0.0001 for uninf versus infected and ##p<0.01; ####p < 0.0001 for DMSO versus LEF treatment by means of two-way ANOVA analysis. (**B**) Total abundance of carbamoyl aspartic acid and dihydroorotic acid in samples treated as in (A). Data are mean ±SEM of n=3 independent cultures, and are normalized by cell number, ****p<0.0001 for uninfected versus infected and ####p < 0.0001 for DMSO versus LEF treatment by means of two-way ANOVA analysis. (**C**) Schematic of bifunctional dihydrofolate reductase thymidylate synthase (DHFR-TS) of *Toxoplasma gondii*.

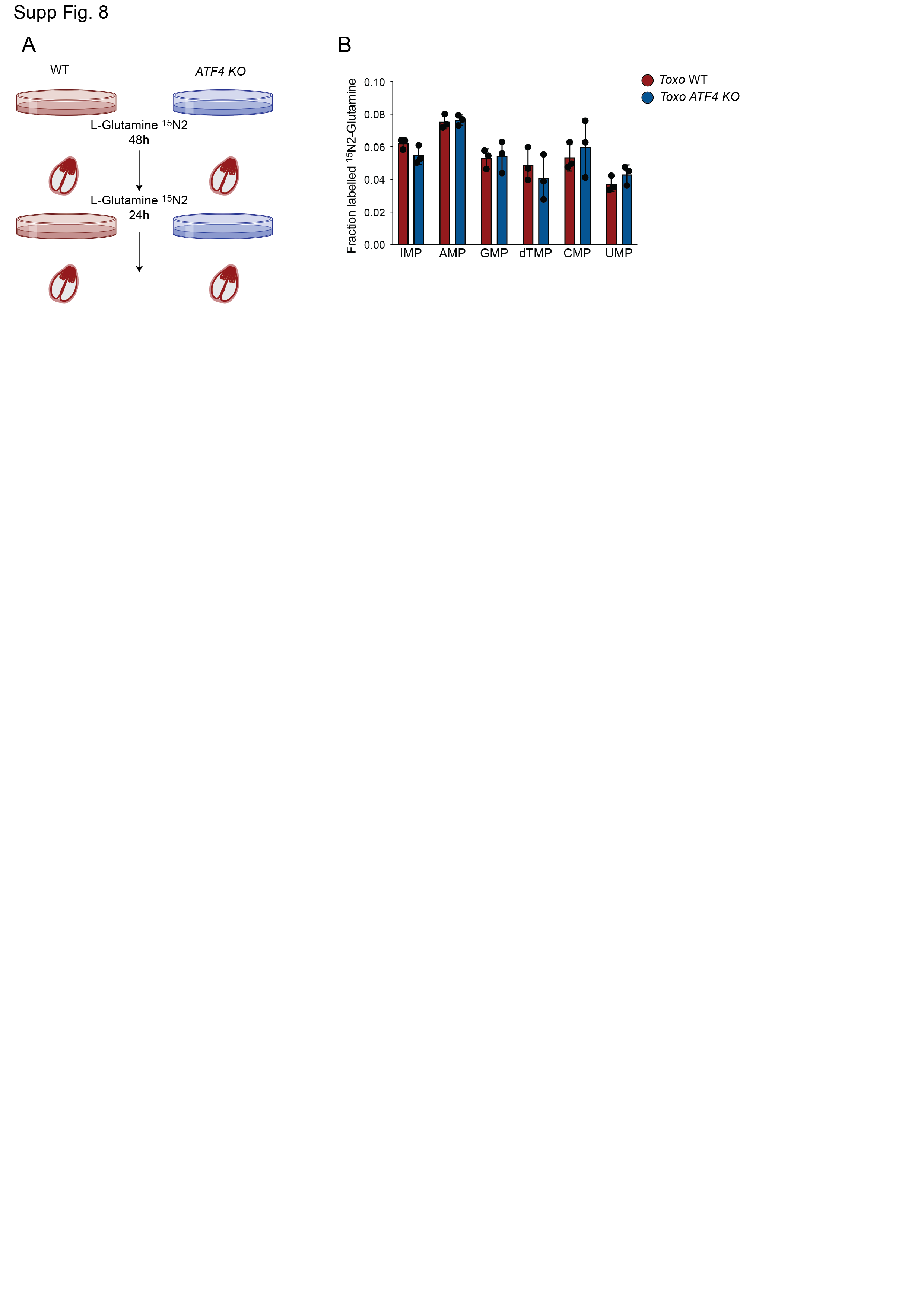
Fig. S8. Loss of ATF4 does not promote nucleotide acquisition by *Toxoplasma*. (A) Workflow for assessing *Toxoplasma* nucleotide acquisition from host cells. WT and *ATF4* KO ES-2 cells were culture with 2 mM L-glutamine^15^N_2_ for 48 h. Cells were then infected with *Toxoplasma* in the absence of the isotopologue L-glutamine^15^N_2_. After 24h the *Toxoplasma*-enriched fraction was pelleted for metabolite extraction. (B) Fraction enrichment of glutamine-derived ^15^N nucleotides of *Toxoplasma*-purified extracts from WT and *ATF4* KO cells as described in (A). Data are ± SD of n=3 independent cultures.

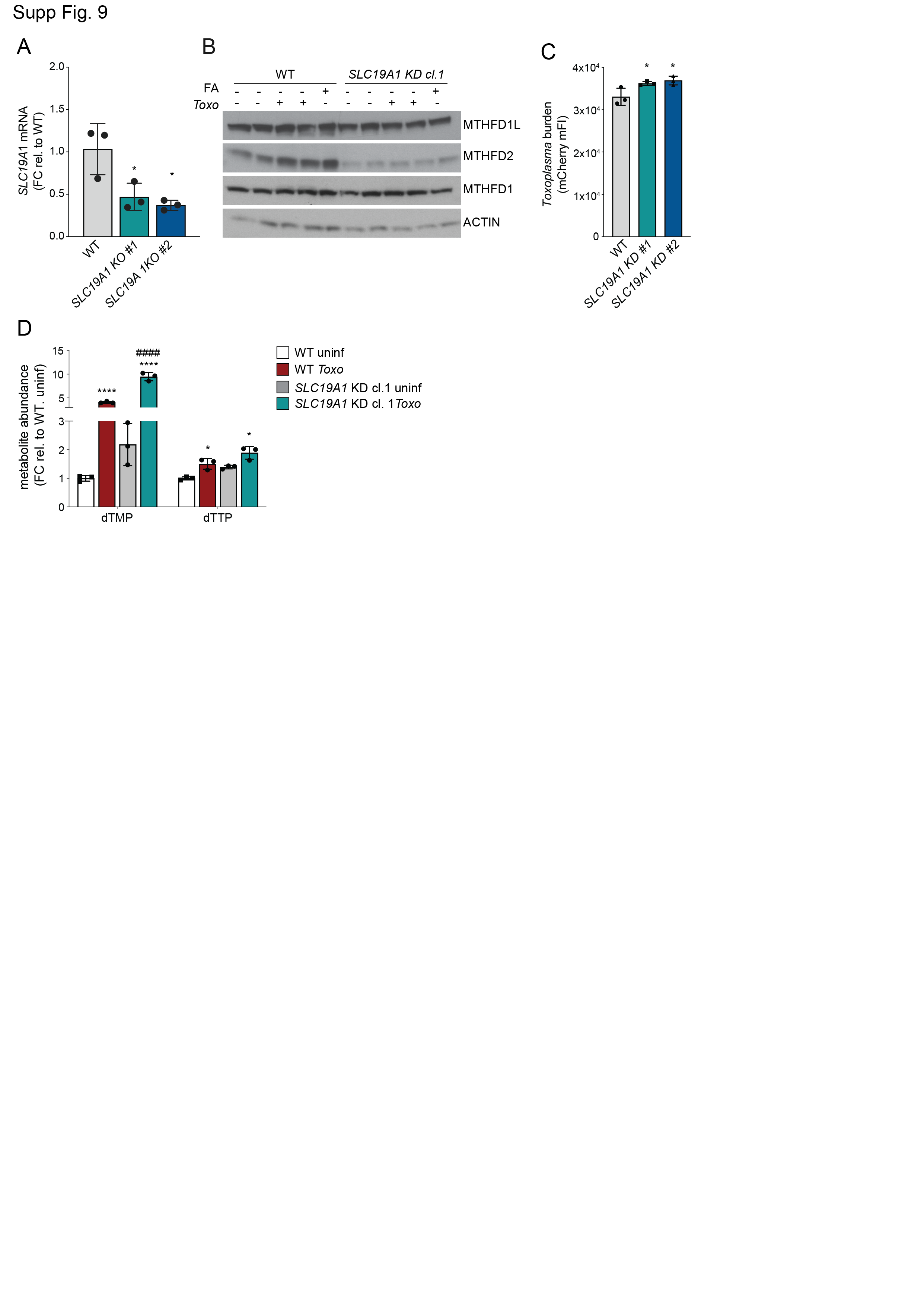

**Fig. S9. Increased parasite proliferation and dTMP levels in a model of mitochondrial 1C-dysfunction.** (**A**) WT and SLC19A KO cells were analyzed by qPCR for *SLC19A1*. Transcripts were normalized to *ACTB*. Data are mean ± SD of n=3 independent cultures. (**B**) WT and *SLC19A1* knockdown (KD) ES-2 cells were cultured for 4 days in RPMI medium lacking folic acid and uninfected, infected with *Toxoplasma* (MOI: 4) cells or supplemented with folic acid (2 μM). After 24h, samples were analyzed by immunoblotting for the indicated antibodies MTHFD1L, ~106 kDa; MTHFD2, ~32 kDa; MTHFD1, ~40 kDa and α-Actin (ACTB), ~35 kDa. (**C**) WT and S*LC19A1* KD ES-2 cells were infected with *Toxoplasma* and analyzed 24 hpi by means of flow cytometry for *Toxoplasma* burden [mCherry median FI]. Data are mean ± SEM of three biological experiments, *p < 0.05 for WT versus *SLC19A1* KO, by means of one-way ANOVA analysis. (**D**) Total abundance of dTMP and dTTP of WT and S*LC19A1* KO ES-2 uninfected and *Toxo*-infected cells. Data are mean ± of SED n=3 independent cultures, and normalized by cell number, *p<0.05; ****p<0.0001 for uninfected versus infected cells and ####p < 0.0001 for WT versus *SLC19A1* KO cells by means of two-way ANOVA analysis.

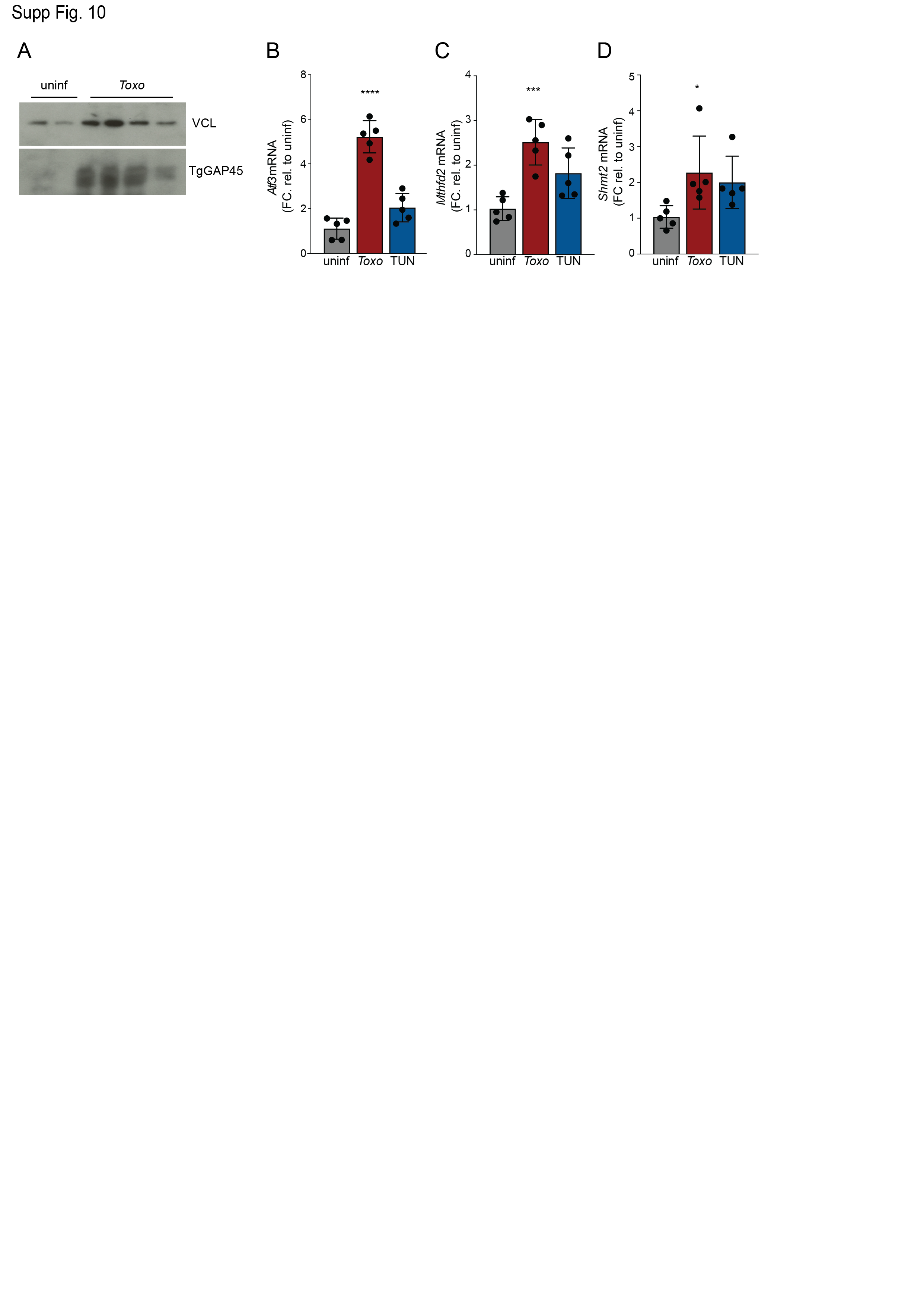
**Fig. S10. *Toxoplasma* infection and tunicamycin treatment induce ATF4 targets in vivo.** (**A**) Immunoblot (IB) analysis of lysates of peritoneal exudate cells (PECs) isolated at 5 dpi from mice that there uninfected or infected with 50 tachyzoites: *Toxoplasma* GAP45 (TgGAP45), ~45 kDa; Vinculin (VCL), ~124 kDa; (**B**-**D**) Mice were injected intraperitoneally with 1x PBS or 50 tachyzoites in 1x PBS or with tunicamycin (1 mg kg^−1^ body weight). At 5 days post injection (dpi) peritoneal exudate cells (PECs) were isolated and analyzed for the indicated ATF4 targets genes by qPCR. Transcripts are normalized to *Hprt* and relative to untreated uninfected PECs mice (n=5). * p<0.05; *** p<0.001; and ****p<0.0001 by means of one-way ANOVA analysis.
